# Matrix metalloproteinase-2 mediates ribosomal RNA transcription by cleaving nucleolar histones

**DOI:** 10.1101/2020.02.21.958280

**Authors:** Mohammad A.M. Ali, Javier A. Garcia-Vilas, Christopher R. Cromwell, Basil P. Hubbard, Michael J. Hendzel, Richard Schulz

## Abstract

Cell proliferation and survival require continuous ribosome biogenesis and protein synthesis. Genes encoding ribosomal RNA (rRNA) are physically located in a specialized substructure within the nucleus known as the nucleolus, which has a central role in the biogenesis of ribosomes. Matrix metalloproteinase-2 (MMP-2) was previously detected in the nucleus. However, its role there is elusive. Herein we report that MMP-2 resides within the nucleolus to regulate rRNA transcription. MMP-2 is enriched at the promoter region of rRNA gene repeats and its inhibition downregulates pre-rRNA transcription. The N-terminal tail of histone H3 is clipped by MMP-2 in the nucleolus and is associated with increased rRNA transcription. Knocking down/out MMP-2 or inhibiting its activity prevents histone H3 cleavage and reduces both rRNA transcription and cell proliferation. In addition to the known extracellular roles of MMP-2 in tumor growth, our data reveal an epigenetic mechanism whereby intranucleolar MMP-2 regulates cell proliferation through histone proteolysis and facilitation of rRNA transcription.

## Introduction

Cell proliferation is a complex and tightly controlled process [1] involving not only DNA replication but also biosynthesis of a large number of proteins. The nucleolus is the most prominent structure in the nucleus where transcription of ribosomal genes encoding ribosomal RNA (rRNA) occurs [2]. However, the function of the nucleolus goes beyond ribosome production as it is also involved in post-transcriptional modification of RNA, cell cycle regulation, stress responses and cell senescence [3].

Multiple cell signals are coordinated through epigenetic regulation, including DNA methylation and post-translational modifications of the N-terminal tails of core histones constituting the nucleosome [4]. The most studied histone post-translational modifications include acetylation, ubiquitylation and methylation of lysines, arginine methylation and serine and threonine phosphorylation [5]. Histone modifications occur on multiple but specific residues. These combinatorial modifications of histones can also act as binding platforms for specific proteins that read particular histone marks, leading to active or silenced genomic regions.

While the roles of many N-terminal tail histone post-translational modifications in gene expression are well established [4], the significance and mechanisms of histone tail proteolysis are less clear. Chromatin modification through the proteolytic cleavage of histones is a rapidly developing topic in epigenetics [6]. Cathepsin L was the first identified protease involved in epigenetic regulation by histone proteolysis in mouse cells [7]. However, the identity of functionally related yeast and human protease(s) has not been fully elucidated [8, 9]. Functionally, histone N-termini are not required for the assembly of the nucleosome per se but are critical in the formation of more compact chromatin structures beyond the 10 nm linear nucleosome structure [10]. Thus, the consequence of histone N-terminal proteolytic removal may be to promote an activated chromatin state that is resistant to chromatin compaction [11–15].

Matrix metalloproteinase-2 (MMP-2) was first identified as a ubiquitously expressed protease which targets extracellular matrix proteins (also named gelatinase A), but is now recognized through its proteolytic activities to be involved in various intracellular processes, both physiological and pathological [16]. MMP-2 is localized in specific cellular compartments and organelles including the nucleus [17], the sarcomere [18], caveolae [19], mitochondria and the mitochondria-associated membrane [20]. The intracellular roles of MMPs are still incompletely understood. The signal sequence of 72 kDa MMP-2 inefficiently targets it to the secretory pathway and approximately half of nascent MMP-2 remains intracellular [21]. Several intracellular targets of MMP-2 have been discovered including titin [22], troponin I [18], α-actinin [23] and glycogen synthase kinase-3β [24]. Activation of intracellular MMP-2 has been linked to pathological conditions such as oxidative stress injuries to the heart [22] and brain [25] as well as tumor growth [26]. MMP-9, a related MMP also found in the nucleus [17] was recently found to epigenetically regulate mouse osteoclast differentiation through cleavage of histone H3 [27].

MMP-2 was the first MMP to be found inside the nucleus [17], yet its biological functions in that organelle remain elusive. We report here that nuclear MMP-2 is enriched at the sites of transcription within the nucleolus. We provide evidence that intracellular MMP-2 co-localizes with nucleolus specific markers and is found in purified nucleolar fractions. Further, we show that MMP-2 is bound to chromatin within the promoter of the ribosomal RNA (rRNA) gene, and that it is directly involved in rRNA gene transcription. We provide data showing that MMP-2 is able to clip the N-terminal tail of histone H3 within the nucleolus, which would reduce the capacity of the chromatin to compact. Finally, we show that silencing or knocking-out MMP-2 inhibits pre-rRNA transcription as well as cell proliferation. The involvement of MMP-2 in ribosomal gene expression sheds light on a novel role this protease may play in oncogenesis.

## Results

### MMP-2 co-localizes with nucleolar markers

MMP-2 has been located in the nucleus of cardiomyocytes [17, 28], endothelial cells [29] and neurons [25]. Closer inspection of our previous immunogold electron microscopy and immunofluorescence data [17, 28] suggested to us that MMP-2 might be present within the nucleolus. Consequently, we wanted to more carefully examine the localization of nuclear MMP-2 in order to gain insight into potential MMP-2 nuclear functions.

Immunofluorescence of MMP-2 revealed staining throughout the cell (Figure 1A). In addition to the expected cytoplasmic staining, there was a clear abundance of signal in the nucleus. Within the nucleus, there were abundant small nuclear foci of MMP-2 staining scattered throughout the nucleoplasm but several foci stood out as brighter and slightly larger (encircled in Fig. 1A). This pattern of MMP-2 staining is distinct from most RNA polymerase II transcription factors whose nucleoplasmic staining reveals an exclusion zone corresponding to the nucleolus [30]. Since MMP-2 staining was found in chromatin-depleted regions of the nucleus, we tested for colocalization with a nucleolar marker, fibrillarin, which localizes to the fibrillar center of the nucleolus. We found that the bright MMP-2 foci co-localized with fibrillarin (Fig. 1A). We confirmed the co-localization of MMP-2 with fibrillarin in several other human cancer cell lines including MCF7, PC-3, and U-373 MG cells (Supplementary Fig. S1).

**Figure 1.**
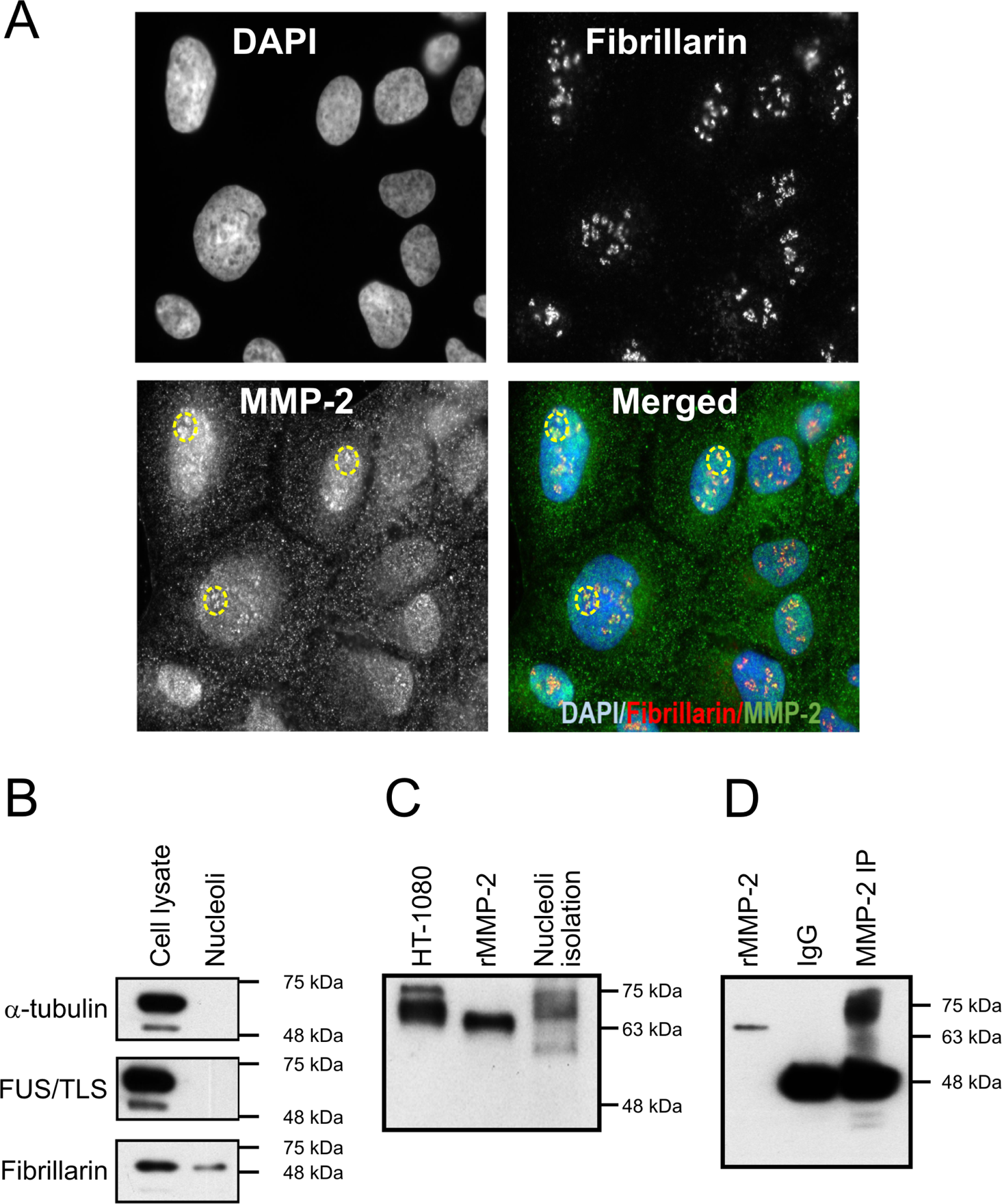
MMP-2 is present in the nucleolus. **A.** Localization of MMP-2 to fibrillarin within nucleoli of human U2-OS cells. Punctate nucleolar staining of fibrillarin (upper right). Examples of MMP-2 foci in nucleoli are encircled (lower left). In the merged image (lower right), MMP-2 is co-localized with fibrillarin; MMP-2 (green), fibrillarin (red) and DAPI (blue). Shown are representative images from n = 4 independent experiments. **B**. MMP-2 is present in purified nucleoli fractions of U2-OS cells. Nucleoli purity was confirmed by the absence of α-tubulin (cytoplasm marker) and FUS/TLS (nuclear marker), and the presence of fibrillarin (specific nucleoli marker). Shown are representative results of n = 7 independent experiments. **C**. MMP-2 presence in purified nucleoli as determined by Western blot. HT-1080 and recombinant human MMP-2 (rMMP-2) were used as positive controls, n = 3 independent experiments. **D.** Western blot of MMP-2 immunoprecipitated from isolated nucleolar lysates. Shown are rMMP-2 standard and immunoprecipitates using either IgG isotype control, or anti-MMP-2 antibody; n = 3 independent experiments.

To validate the presence of MMP-2 within the nucleolus, we isolated highly purified nucleoli from U2-OS cells. We confirmed their purity by the absence of α-tubulin (cytoplasmic marker), FUS/TLS (nuclear marker), and by the presence of fibrillarin in Western blots (Fig. 1B). A Western blot for MMP-2 in the nucleoli preparation showed a band of approximately 72 kDa, in accordance with recombinant human MMP-2 and conditioned media from human HT-1080 cells (commonly used as an MMP-2 standard) (Fig. 1C). MMP-2 was detected in immunoprecipitates from purified nucleoli as a prominent 72 kDa band, which was absent in IgG control immunoprecipitate (Fig. 1D). These data strongly suggest the presence of MMP-2 within the nucleoli of human cancer cell lines.

### MMP-2 localizes to the sites of active transcription within the nucleolus and binds ribosomal DNA

To investigate a potential role of MMP-2 within the nucleolus, we performed pulse-labeling experiments with 5-fluorouridine to detect nascent rRNA synthesis. A 10 min pulse of 5-fluorouridine in U2-OS cells revealed a classical pattern of clusters of small discrete foci within the larger nucleolar body (Fig. 2A). Double staining for 5-fluorouridine and MMP-2 revealed a tight association between MMP-2 and rRNA transcription [Fig. 2A and 2B].

**Figure 2.**
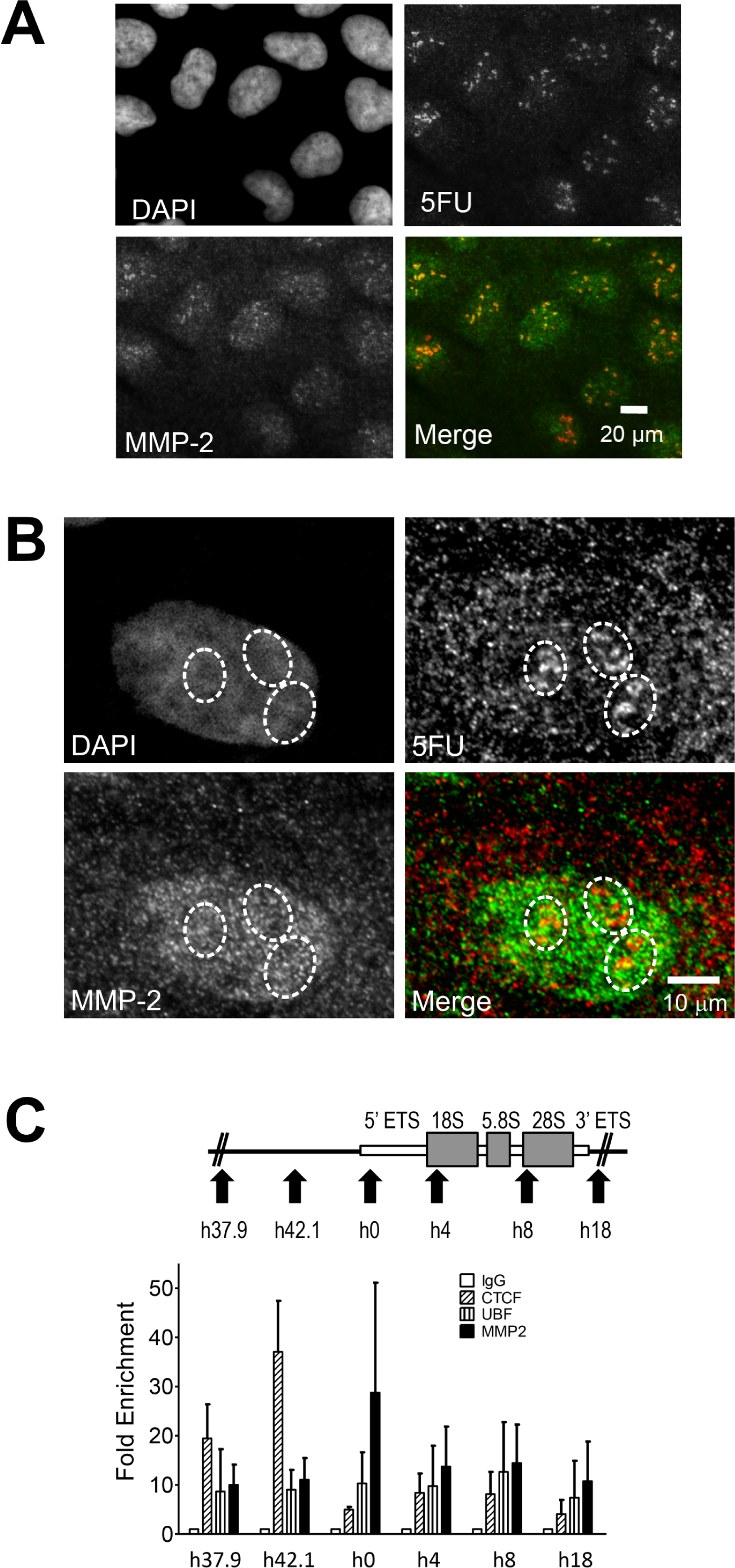
MMP-2 is present in the nucleoli at transcriptionally active sites. **A** and **B**. Localization of MMP-2 to active transcription sites within nucleoli. Human U2-OS cells were pulsed with 5-fluorouridine (5FU) and stained with MMP-2 and BrdU antibodies. Merged image shows co-localization of MMP-2 (green) with 5FU (red) within nucleoli at low 40x (**A**) and high 60x (**B**) magnification as denoted (scale bars as indicated). **B.** Sites of double staining for 5FU and MMP-2 (yellow) are encircled and show presence of MMP-2 within active transcription sites in nucleoli, n = 4 independent experiments. **C**. Upper panel: diagram (not to scale) of rDNA gene showing the location of the primer sets used in nucleolar ChIP-qPCR assay. Lower panel: nucleolar ChIP shows that MMP-2 is highly enriched at the promoter region of the rDNA gene (h0 locus). CTCF and UBF were used as controls. CTCF is enriched in the spacer promoter h37.9 and h42.1 loci, whereas UBF is enriched across the rDNA gene, n = 3 independent experiments.

Given the apparent localization of MMP-2 within the nucleolus to sites of active transcription, we next sought to determine whether MMP-2 associates with nucleolar chromatin. Consequently, we performed nucleolar chromatin immunoprecipitation (ChIP) assays as previously described [31], followed by qPCR using primer sets mapping different loci (h0, h4, h8, h18, h37.9 or h42.1) across the rDNA gene (Fig. 2C). Antibodies against both CCCTC-binding factor (CTCF) and upstream binding transcription factor (UBF), known factors involved in rDNA gene transcription, were used as positive controls. As previously reported, CTCF was enriched mainly to the spacer promoter regions h37.9 and h42.1 [32], whereas UBF showed enrichment across the rDNA gene [33]. Interestingly, we found that MMP-2 binds to different regions of the rDNA gene and is mainly enriched at h0 locus of the rDNA gene promoter (Fig. 2C). This is consistent with a role for MMP-2 in regulating rRNA transcription initiation.

### MMP-2 clips histones at their N-termini

Given that the related protease, MMP-9 was shown to positively regulate RNA pol II transcription through proteolytic removal of histone H3 N-terminus [27] we tested whether MMP-2 can proteolyze nucleosomal histones; (H2A, H2B, H4 and H3 isoforms). Incubation with MMP-2 resulted in concentration-dependent proteolysis of all tested recombinant histones with histone H3 isoforms showing the highest susceptibility (Fig. 3). Inhibition of MMP-2 activity with the MMP-2 preferring inhibitor ARP-100 (10 µM) prevented histone proteolysis.

**Figure 3.**
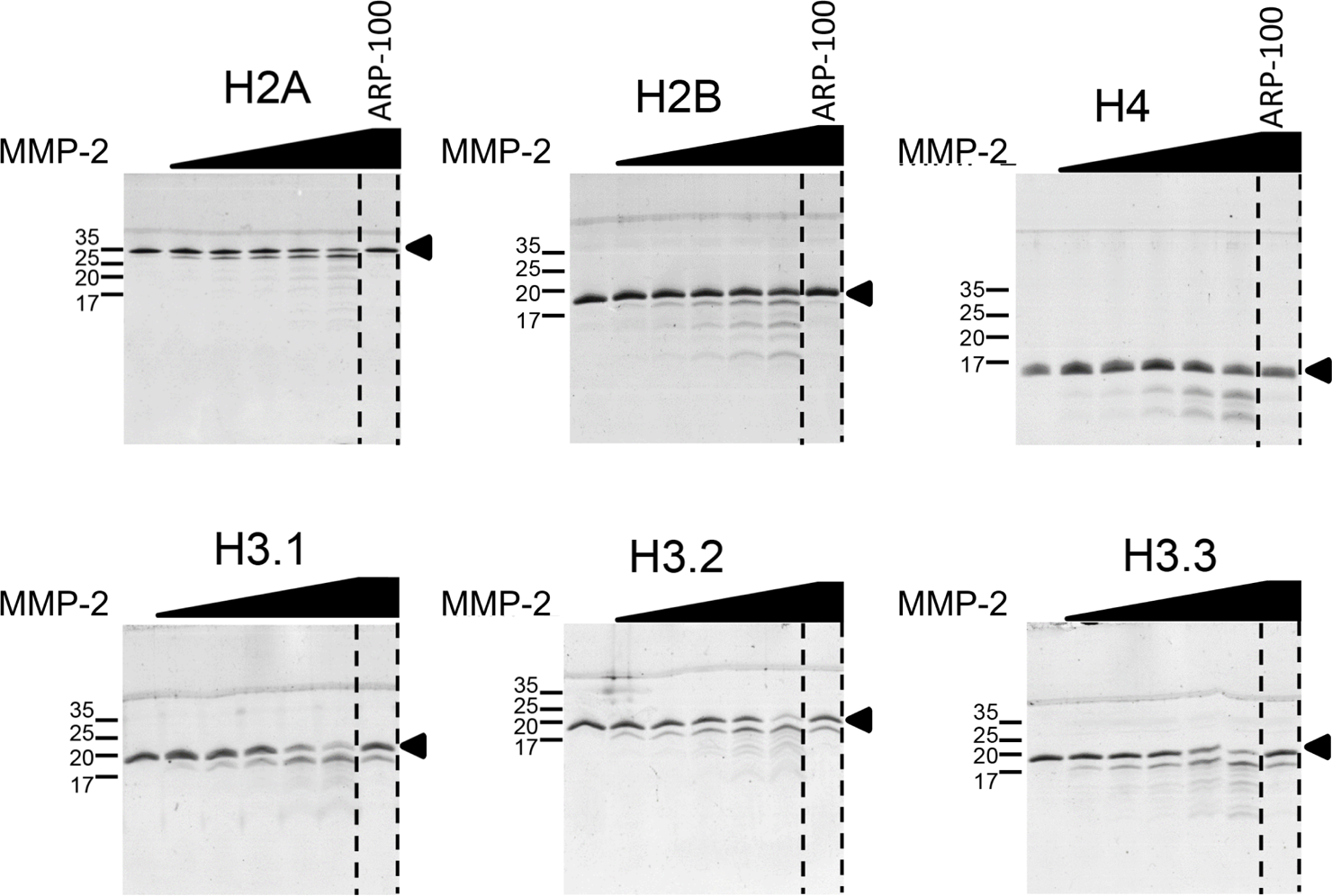
MMP-2 degrades histones *in vitro*. Representative images of proteolysis assays (1 h, 37°C) with recombinant human histones and MMP-2, at increasing MMP-2: histone molar ratios (1:1000 to 1:50). Coomassie blue stained gels are shown in each panel. ARP-100: MMP-2 preferring inhibitor (10 μM). Uncleaved histone bands are indicated with arrowheads. Molecular weight markers shown at left, n = 3 independent experiments.

We then focused on MMP-2 induced proteolysis of histone H3 due to its well-known role in epigenetic regulation of gene expression [34]. After carrying out *in vitro* proteolysis by MMP-2 with different histone H3 isoforms, we did Western blotting using specific antibodies against histone H3 N-terminal or C-terminal domains (Fig. 4A). We observed histone H3 degradation products only when using C-terminal antibodies, indicating that MMP-2 mediated proteolysis occurs at their N-termini.

**Figure 4.**
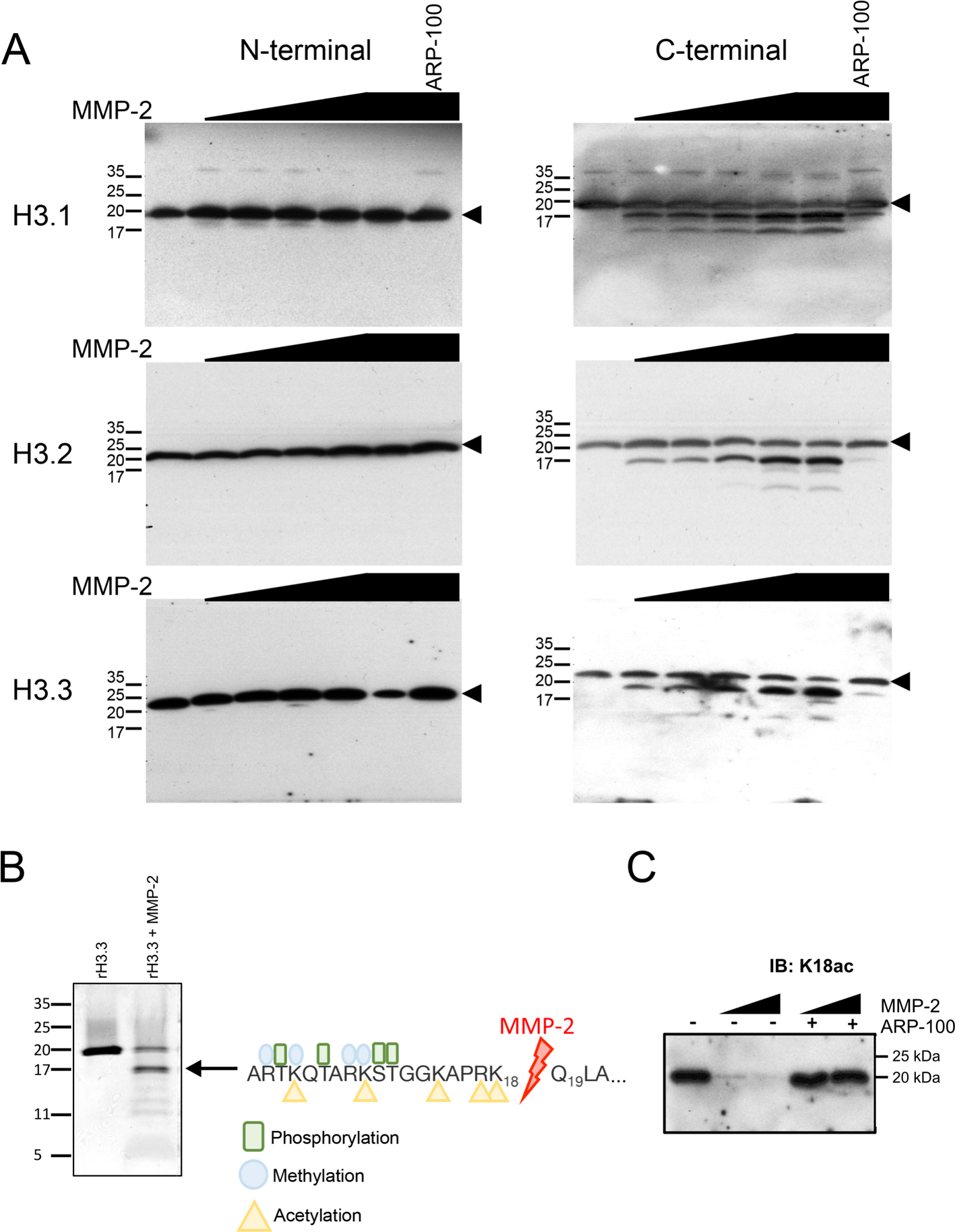
MMP-2 cleaves the N-terminus of histone H3. **A.** Western blots using antibodies against N-terminal (left) and C-terminal (right) specific epitopes in recombinant human histone H3 isoforms after incubation (1 h, 37°C), with increasing molar ratios of recombinant MMP-2 to histone (1:1000 to 1:50). In the right lane of each panel, using the highest molar ratio, 10 µM ARP-100 (MMP-2 preferring inhibitor) was added to confirm MMP-2-dependent histone clipping. Arrowheads show the uncleaved histone band. Molecular weight markers are shown at left, n = 3 independent experiments. **B.** Left: In vitro degradation of recombinant human histone H3.3 (rH3.3) by MMP-2 at 1:150 (MMP-2:histone) molar ratio for 1 h at 37°C. Arrow shows the abundant ≈17 kDa cleavage product of rH3.3 by MMP-2. Right: Edman degradation sequencing of this band shows an MMP-2 cleavage site in rH3.3 between K18 and Q19 residues, n = 3 independent experiments. **C.** Histones purified from isolated U2-OS nucleoli were incubated with MMP-2 (1 h, 37°C) with or without 10 µM ARP-100. Immunoblot using anti-H3 K18ac does not reveal the 17 kDa cleavage product, verifying that MMP-2 cleavage of histone H3.3 occurs at its N-terminus, n = 3 independent experiments.

We next performed N-terminal sequencing of recombinant histone H3.3 that had been incubated at 37°C for 1 h with MMP-2. Recombinant histone was used to avoid interference in protein sequencing due to posttranslational modifications. Although we observed at least three degradation bands of histone H3.3 by immunoblot (Fig. 4B), the most prominent proteolytic product migrating at ∼17 kDa was also seen with shorter incubation times down to 15 min (data not shown). This 17 kDa band was isolated and sequenced and the results showed that the initial cleavage site is located between the K18 and Q19 residues, G13KAPRK18-Q19LATKA (Fig. 4B and see the Supplement for mass spectrometry data). Using *in silico* analysis with CleavPredict [35], we also identified other potential MMP-2 cleavage sites at amino acid positions R41, T46, A48, E51, E74, D92 and R130.

We next determined whether exogenous MMP-2 cleaves native histone H3 isolated from nucleoli near its N-terminus. We used histone H3 antibodies specific against posttranslational modifications of the amino acids nearest to the predicted K18-Q19 MMP-2 cleavage site. Purified nucleoli were isolated from U2-OS cells, and then nucleolar histones were isolated and incubated with MMP-2 for 4 h at 37°C. Figure 4C shows that there was no detectable histone H3 cleavage product using anti-H3 K18_ac_, in contrast to the approximately 17 kDa band revealing a histone H3 degradation product detected using C-terminal anti-H3 (Fig. 4B). This histone H3 cleavage product was also detected using anti-H3 K23ac, anti-H3 K27m_3_ or anti-H3 K36m_3_ (data not shown). Our results show that an important MMP-2 cleavage site on both recombinant and nucleolar histone H3 is located at its N-terminus between the K18 and Q19 residues.

### Nucleolar MMP-2 cleaves histone H3 in nucleoli

To explore whether endogenous MMP-2 is able to cleave histone H3 in situ within the nucleolus, we first tested for a potential interaction between them. MMP-2 was immunoprecipitated from purified nucleoli from U2-OS cells using anti-MMP-2 (ab92536). Western blots showed that histone H3 co-immunoprecipitates with MMP-2 within nucleoli (Fig. 5A). To evaluate whether MMP-2 within the intact nucleolus has proteolytic activity, we carried out degradation assays using isolated purified nucleoli. Nucleolar preparations were incubated for 4 h at 4°C or 37°C, electrophoresed, and immunoblotted with anti-histone H3 C-terminal antibody. Our results showed histone H3 is clipped in the purified nucleolar fractions incubated at 37°C but not 4°C. Importantly the MMP-2 inhibitor ARP-100 prevented histone H3 clipping (Fig. 5B). This suggested the presence of MMP-2 proteolytic activity in purified nucleolar preparations.

**Figure 5.**
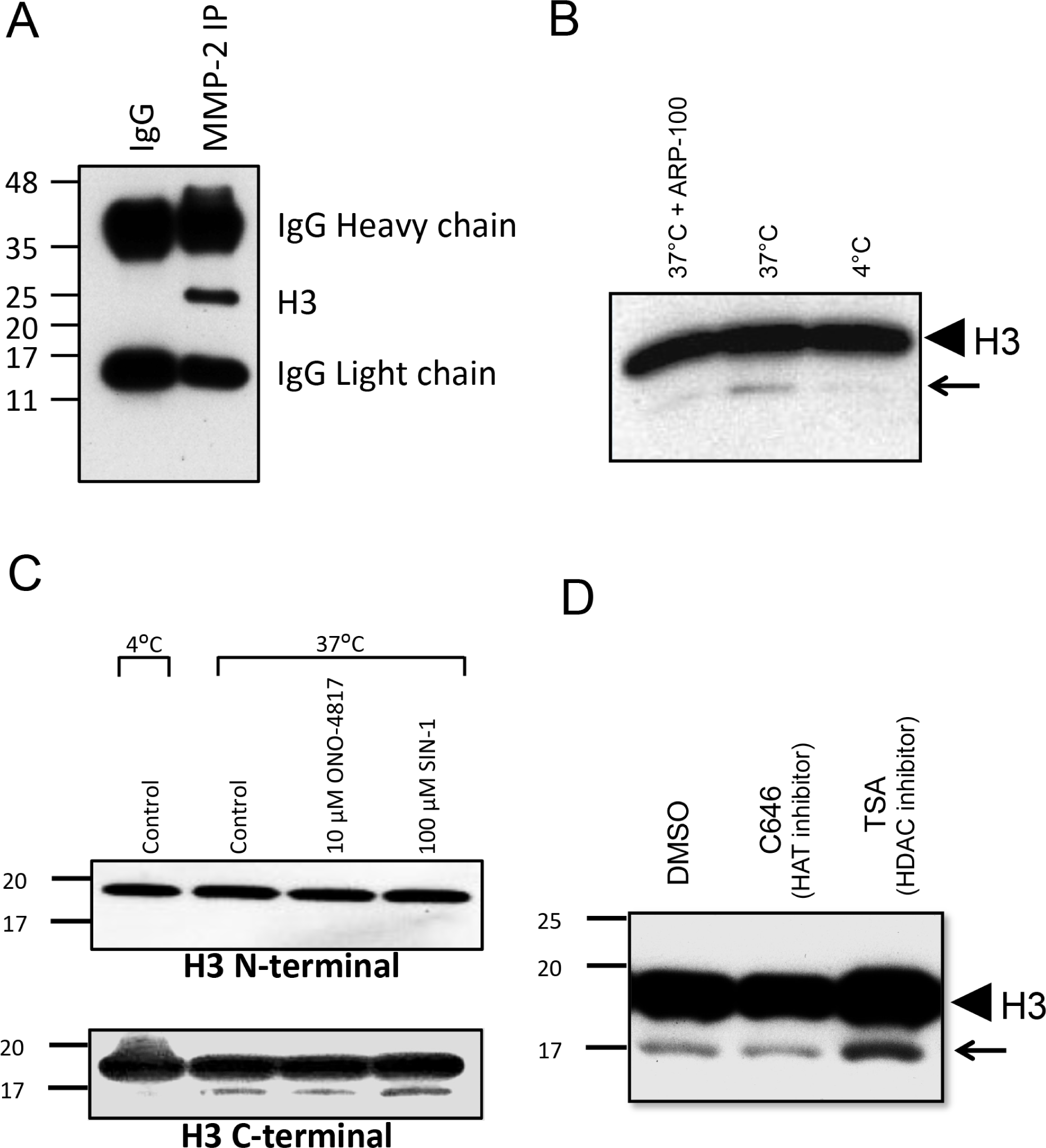
Histone H3 is an MMP-2 substrate in the nucleolus. **A**. Histone H3 co-immunoprecipitates with MMP-2 in purified nucleoli isolated from U2-OS cells. Immunoprecipitation was carried out with anti-MMP-2 or control IgG antibodies. Western blot was performed using anti-H3 C-terminal antibody (n = 3 independent experiments). **B**. Histone H3 proteolysis by endogenous MMP-2 in nucleoli. Lysed nucleoli isolated from U2-OS cells were incubated for 4 h with or without 10 µM ARP-100 at 37°C or at 4°C. Western blot using anti-H3 C-terminal antibody was performed. Arrowhead shows histone H3 and arrow indicates degradation product of histone H3, n = 3 independent experiments. **C**. U2-OS cells were treated with DMSO (control), ONO-4817 or SIN-1 for 3 days. Nucleoli were then isolated and incubated for 1 h at 37°C. Control samples were incubated at 4°C or at 37°C to determine basal proteolytic activity of endogenous MMP-2 on histone H3 in situ. Western blots were done using anti-H3 N-terminal (upper panel) or anti-H3 C-terminal (lower panel) antibodies. **D**. Effect of histone acetylation on its cleavage by nucleolar MMP-2. U2-OS cells were treated with ONO-4817 for 3 days, followed by DMSO, C646 (histone acetyltransferase inhibitor) or trichostatin-A (TSA, histone deacetylase inhibitor) for 1 day. Anti-H3 K27ac antibody was used to determine whether MMP-2 cleaves histone H3 in an acetylation-dependent manner, n= 3 independent experiments.

In order to test that the N-terminus of histone H3 in situ is the site of cleavage by endogenous MMP-2, U2-OS cells were treated either with a cell-permeable MMP inhibitor that preferentially inhibits MMP-2, ONO-4817 (10 μM), or with a peroxynitrite donor, SIN-1 (100 μM), to activate endogenous MMP-2 by oxidative stress [36]. Neither compound was cytotoxic at the selected concentrations (Supplementary Fig. S2). The cells were treated for 3 days, given that the half-life of histone H3.3 turnover is about 3.5 days [37]. As shown in Figure 5C, nucleolar MMP-2 was able to cleave the N-terminus of histone H3 in nucleoli to a 17 kDa fragment detected using a histone H3 C-terminal antibody. This cleavage was decreased by ONO-4817 and increased with SIN-1 treatment.

As MMP-2 cleavage of histone H3 occurred at a well-recognized acetylation site (K18) [5], we tested whether acetylation of histone H3 affects its cleavage by MMP-2. To this end, U2-OS cells were first treated with ONO-4817 for 3 days, and then treated for an additional 24 h with C646, a histone acetyltransferase inhibitor, or with trichostatin-A, a histone deacetylase inhibitor. C646 enhances the proportion of deacetylated histones in U2-OS cells, whereas trichostatin-A enhances histone acetylation. After these treatments, the purified nucleoli were isolated and an immunoblot against histone H3 C-terminus was performed on the nucleolar extract. We observed enhanced histone H3 cleavage to a product migrating at 17 kDa in trichostatin-A-treated cells and attenuated histone H3 cleavage in C646-treated cells (Fig. 5D). Taken together, our results confirm that MMP-2 from nucleoli is proteolytically active and cleaves the N-terminus of histone H3 in an acetylation dependent manner.

### MMP-2 mediates rRNA transcription and cancer cell proliferation

To test whether there is a functional role of MMP-2 at transcription sites in the nucleolus, we silenced MMP-2 in U2-OS cells using two specific siRNAs against MMP-2, which effectively diminished its protein level and activity (Fig. 6A) as well as its mRNA level (Fig. 6B). Importantly, the siRNAs did not affect MMP-9 activity (Fig. 6A). MMP-2 knockdown reduced the synthesis of pre-rRNA by approximately 36% (siMMP-2_A_) and 30% (siMMP-2_B_) compared to siRNA control (Fig. 6C). Likewise, synthesis of pre-rRNA was significantly diminished to a similar extent when cells were treated with either of the two MMP-2 inhibitors (ARP-100 or ONO-4817). As expected, the RNA polymerase inhibitor actinomycin D, used as a positive control, abolished pre-rRNA expression (Fig. 6D).

**Figure 6.**
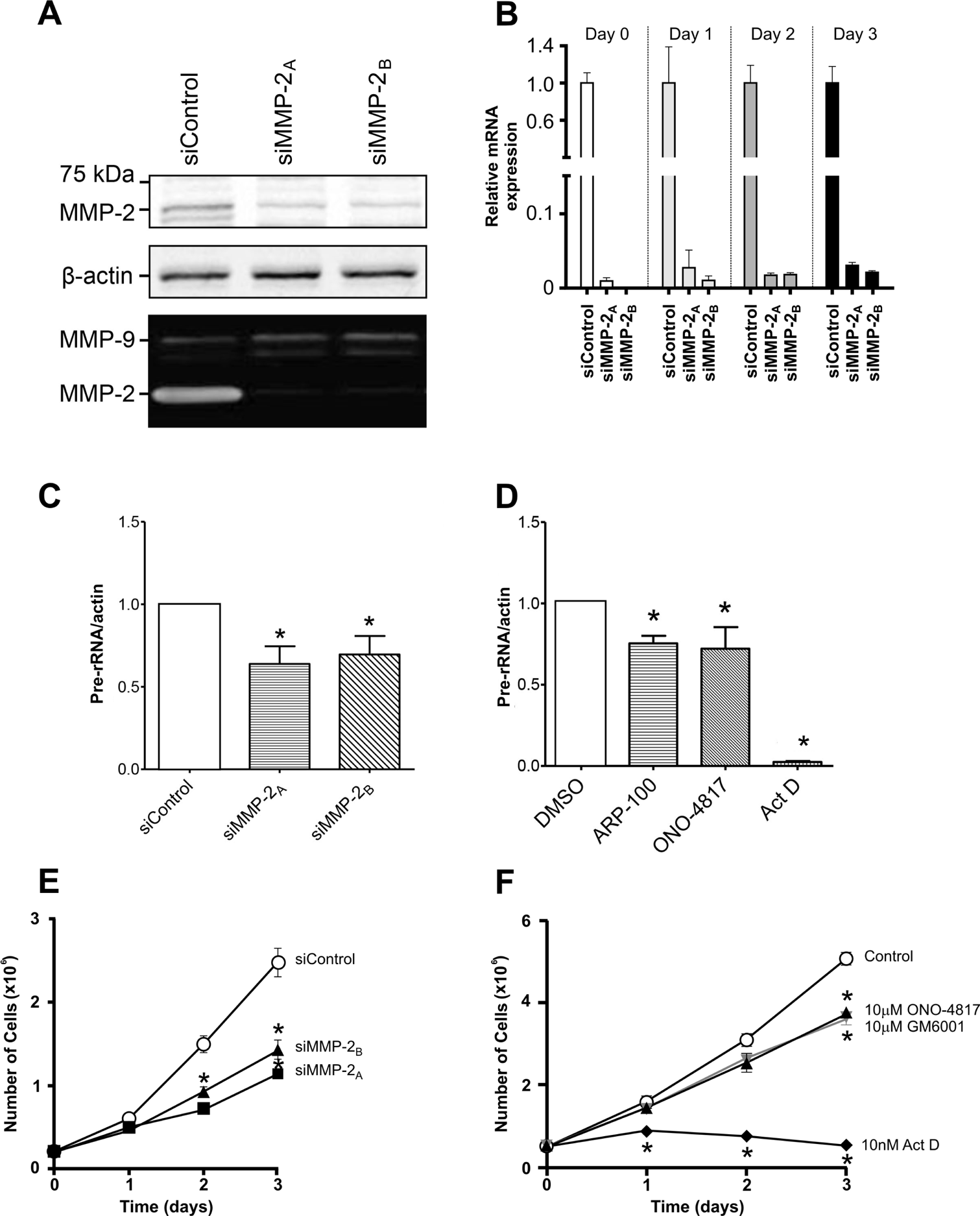
Silencing of MMP-2 affects cell proliferation and pre-rRNA transcription. **A**. MMP-2 knockdown in U2-OS using two different siRNAs (siMMP-2_A_ or siMMP-2_B_) in comparison to negative control (siControl). Upper panel: Western blot of MMP-2 protein levels in cell lysates. Lower panel: gelatin zymography showing MMP-2 activity in cell lysates. **B**. RNA expression of MMP-2 by qPCR, normalized to HPRT1, in MMP-2 knockdown cells using two different siRNAs (siMMP-2_A_ or siMMP-2_B_), relative to siControl (set as 1). **C.** MMP-2 knockdown with siRNAs (siMMP-2_A_ or siMMP-2_B_) reduces pre-rRNA levels in total RNAs measured by qPCR. **D.** Inhibiting MMP-2 activity with ARP-100 or ONO-4817 reduces pre-rRNA levels in total RNAs measured by qPCR. Actinomycin D (Act D) was used as a positive control. **E.** Knocking down MMP-2 with two different siRNAs (siMMP-2_A_ or siMMP-2_B_) significantly inhibits cell proliferation in comparison to siControl. **F.** Inhibiting MMP-2 activity with distinct MMP inhibitors (ONO-4817: MMP-2 preferring, or GM6001: pan MMP inhibitor) reduces cell proliferation in comparison to DMSO control. Actinomycin D (Act D) was used as a positive control, n=3 independent experiments for panels A-F, *p<0.05 two-way ANOVA.

We then measured U2-OS cell proliferation over 3 days in both silenced and control cells. Figure 6E shows that silencing MMP-2 using siRNAs results in a slower cell proliferation rate than cells transfected with control siRNA. We also addressed the biological role of MMP-2 using cells treated with MMP inhibitors (MMP-2 preferring: ONO-4817; pan-MMP: GM6001) at non-cytotoxic concentrations (Supplementary Fig. S2). Cell proliferation assays revealed that either ONO-4817 or GM6001 significantly decreased the rate of cell proliferation compared to control (Fig. 6F).

To further confirm the role of MMP-2 in cell proliferation, we used the CRISPR/Cas9 system to knockout the MMP-2 gene in U2-OS cells. The mRNA, protein level and activity of MMP-2 were abolished in the CRISPR/Cas9 MMP-2 knockout cells (Fig. 7A). Moreover, pre-rRNA levels were significantly reduced in MMP-2 deficient cells (Fig. 7B). Most importantly, the proliferation rate of MMP-2 deficient U2-OS cells was greatly diminished (Fig. 7C). To confirm whether the phenotype of MMP-2 deficient U2-OS cells is indeed caused by depleting MMP-2, MMP-2 was reintroduced into MMP-2 deficient cells by transient transfection. This partially rescued both pre-rRNA levels (Fig. 7B) and the defect in cell proliferation in MMP-2 knockout cell lines (Fig. 7C). In summary, these data indicate that knockdown or knockout of MMP-2 or inhibition of its activity reduces rRNA transcription and inhibits the proliferation of U2-OS cells.

**Figure 7.**
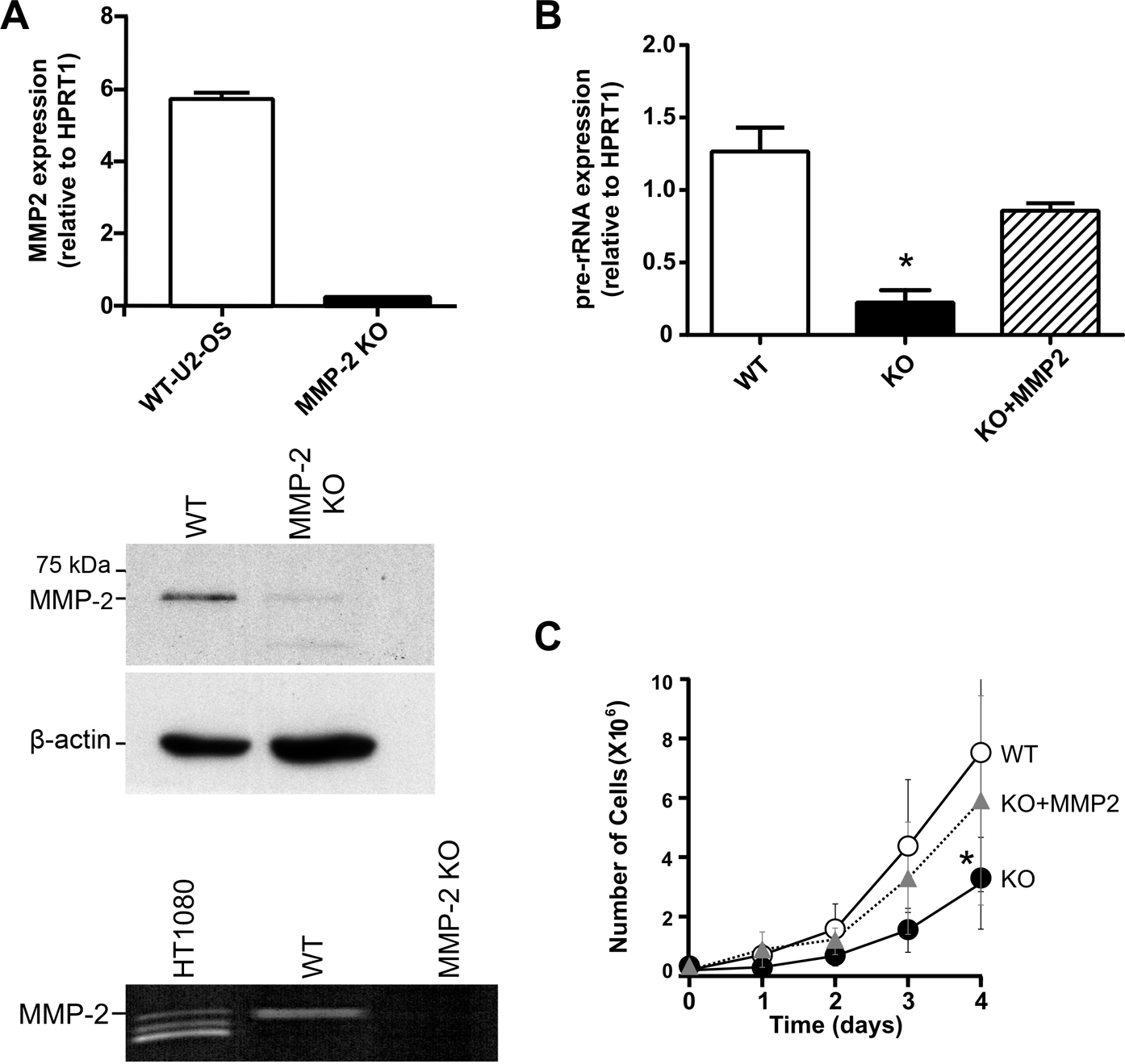
MMP-2 knockout by CRISPR/Cas9 reduces pre-rRNA levels and cell proliferation. **A.** Validation of CRISPR/Cas9 MMP-2 knockout U2-OS cells. MMP-2 mRNA in total RNAs by qPCR (upper panel), MMP-2 protein in cell lysates by Western blot (middle panel) and MMP-2 activity in cell lysates by gelatin zymography (lower panel) show reduced MMP-2 in MMP-2 knockout (KO) cells in comparison to wild type (WT) cells. **B.** pre-rRNA levels measured by qPCR in wild type (WT), MMP-2 knockout (KO) and MMP-2 knockout transfected with MMP-2 construct (KO + MMP-2) U2-OS cells, n=3 independent experiments. **C.** Cell proliferation assay of wild type (WT), MMP-2 knockout (KO) and MMP-2 knockout transfected with MMP-2 construct (KO + MMP-2) U2-OS cells, n=3 independent experiments, *p<0.05 two-way ANOVA.

## Discussion

In this study we defined the subnuclear localization and biological role of MMP-2. MMP-2 had previously been reported to have a nuclear pool [17], but its location within the nucleus and its nuclear functions have not been determined. While MMP-2 is found in a large number of small foci scattered throughout the nucleoplasm, we also observed that it enriches in a specific sub-compartment within the nucleolus, the site of rRNA transcription known as the fibrillar center. Consistent with this, we found that MMP-2 binds the rDNA promoter region and we provide evidence that MMP-2 regulates cancer cell proliferation, at least in part, through rRNA transcription. Furthermore, MMP-2 cleaves the N-terminus tail of histone H3 between the K18 and Q19 residues in nucleolar chromatin. Histone H3 clipping within the nucleolus provides a molecular mechanism by which MMP-2 could epigenetically alter rRNA transcription and thus cell proliferation.

Using unbiased degradomics approaches, several nuclear proteins have been suggested to be potential MMP-2 substrates, including histones [38]. Amongst potential targets of MMP-2, histones are of great interest. Post-translational modifications of histones through N-terminal tail clipping affects nucleosome stability by removing repressive signals to thus induce gene expression [8]. The N-terminus of histone H3, the longest N-tail of core histones, protrudes from the nucleosome at the entry point of the DNA [39]. This tail is a very important substrate for epigenetic regulation through post-translational modifications [40]. Santos-Rosa *et al.* reported that, in yeast, histone H3 N-tail clipping (a cleavage between Q19 and L20) was directly associated with gene promoters following the induction of transcription, and that abrogation of this tail clipping impaired gene expression [8]. The cleavage at Q19 removes 3 arginines and 4 lysines from the N-terminus (Fig. 4B). Apart from eliminating the potential for silencing through K9 methylation, this will reduce the net charge of the N-terminus by as much as +7, which reduces the charge more than what is possible by histone acetylation. This is expected to promote an accessible chromatin state [5, 8, 41].

The nucleolus is divided in three components (fibrillar centre, dense fibrillar component and granular component), each with specific functions and consisting of unique proteins. At the boundary between the dense fibrillar component and fibrillar centre, rRNA transcription takes place [42]. Interestingly, MMP-2 staining within the nucleolus resembled that of nucleolar proteins involved in the regulation of rRNA transcription, including CTCF-L/BORIS [43] and nucleolin [44]. These observations prompted us to test whether MMP-2 binds to rDNA gene repeats and/or regulates rRNA transcription. Indeed, our nucleolar ChIP-qPCR assay mapped MMP-2 to different loci of rDNA gene repeats similar to UBF [33], albeit MMP-2 was more enriched at the gene promoter locus. This indicates that MMP-2 potentially regulates rRNA transcription initiation. Furthermore, our data showed that genetic or pharmacologic inhibition of MMP-2 decreased pre-rRNA transcription. This strongly suggests that MMP-2 activity within the nucleolus facilitates rRNA gene transcription.

Recently MMP-9 was found to epigenetically regulate gene expression during osteoclastogenesis in mice through histone H3 proteolysis [27]. In this study, MMP-9 was also shown to cleave histone H3 between K18 and Q19, which resulted in an upregulation of genes involved in osteoclast differentiation. Interestingly, the cleavage was stimulated by CBP/P300-mediated acetylation of K18. Our data also showed, in cells treated with a histone deacetylase inhibitor, that acetylation of nucleolar histone H3 renders it more susceptible to MMP-2 proteolysis. However, MMP-9 was shown to regulate RNA polymerase II-transcribed genes outside the nucleolus and the authors provided no evidence of a nucleolar association of MMP-9 [27]. To rule out a possible role of MMP-9 in our study, we used two different siRNAs that efficiently knocked down MMP-2 without affecting MMP-9. Thus a subset of MMPs may regulate the transcription of genes involved in cell differentiation and proliferation through a proteolytic mechanism.

We demonstrated that either inhibiting MMP-2 activity, knocking down or knocking out MMP-2 decreased both rRNA transcription and proliferation rate of human U2-OS cells. Interestingly, previous work using targeted knockdown of MMP-2 by siRNA in another human osteosarcoma cell line, SaOS2, or in murine osteoblast MC3T3 cells, resulted in decreased cellular proliferation rates [45]. Moreover, bone cells isolated from MMP-2^-/-^ mice exhibited slower growth rate in comparison to MMP-2^+/+^ cells [45]. Intriguingly, this deficit in cell growth was not recovered by the replacement of MMP-2 in the growth media [45], which clearly suggests that the effect of MMP-2 on cell growth is beyond its role as an extracellular protease. The Lovett group also showed reduced proliferation of rat kidney mesangial cells upon MMP-2 knockdown [46]. These studies, however, did not provide a molecular mechanism by which MMP-2 could enhance cell proliferation. We hereby provide a mechanism which could explain these previous observations.

Collectively, we discovered that MMP-2 resides within the nucleolus where it is bound to the rDNA promoter and associates with histone H3. We propose that nucleolar MMP-2 regulates rRNA transcription and cell proliferation through proteolyzing histone H3 at its N-terminus. This may facilitate rRNA transcription within the nucleolus, consistent with our results showing a requirement for MMP-2 in ribosomal gene transcription. MMP-2 was also found in a large number of small foci distributed throughout the nucleoplasm, where it may regulate RNA polymerase II-transcribed genes in a manner similar to what has been demonstrated for MMP-9. Consequently, it is possible that some contribution to the proliferation defect in the knockout of MMP-2 is mediated through effects on RNA polymerase II-transcribed genes that regulate growth outside of the nucleolus. Our findings, however, highlight a novel role of MMP-2 in ribosomal gene expression processes and expand the inventory of relevant intracellular roles for this ubiquitous protease. Importantly, this provides a completely separate mechanism, distinct from extracellular matrix turnover, whereby MMP-2 can contribute directly to cancer progression. Our study provides an alternative rationale and disease setting for the deployment of MMP-2 inhibitors in the treatment of human tumors.

### Materials and Methods Antibodies and reagents

The following reagents and antibodies were purchased from the indicated sources: Dulbecco’s modified Eagle’s medium (DMEM), Biowhittaker; fetal bovine serum (FBS), Harlan-Seralab; anti-FUS/TLS (ab23439), anti-α-tubulin (ab4074), anti-fibrillarin (ab5821), anti-MMP-2 (ab92536 for WB, IP), anti-H3 C-terminal (ab12079) and anti-H4 C-terminal (ab31827), Abcam; anti-H3 N-terminal (05-499), Upstate Biotechnology; anti-H4 N-terminal (AHP413), Biolegend; anti-H3 K18ac (AR-0103) and anti-H3 K23ac (AR-0104), Upstate Biotechnology; recombinant human histones, New England Biolabs; 72 kDa MMP-2 and anti-MMP-2 (AB-19015, IF), EMD Millipore; anti-CTCF (#2899), Cell Signaling; anti-UBF (sc-13125x), Santa Cruz. Unless otherwise stated, other reagents and chemicals were purchased from Sigma-Aldrich.

### Cell culture

Human osteosarcoma cell line (U2-OS; ATCC-HTB-96), human breast cancer cell line (MCF7; ATCC-HTB-22), human prostatic carcinoma cell line (PC-3; ATCC-CRL-1435), and human glioblastoma cell line (U-373-MG; Sigma-08061901) cells used in this study were maintained at 37°C in a humidified 5% CO_2_ atmosphere in DMEM supplemented with 10% FBS.

### Isolation of nucleoli

Nucleoli from U2-OS cells were isolated as described [47]. Briefly, cells were washed, harvested, and resuspended in nucleoli lysis buffer. Cells were disrupted and cell homogenate was centrifuged at 1,500 x g for 5 min at 4°C. The supernatant was removed and the pellet was resuspended in 500 μL solution-I. The suspension was deposited on 700 μL solution-II, and centrifuged at 2,500 x g for 5 min at 4°C. The supernatant was aspirated and the nuclei-containing pellet was resuspended in 500 μL solution-II. The isolated nuclei were sonicated on ice at 50% amplitude, using 10 s on/off cycles. Sonicated nuclei were then deposited on 700 μL solution-III and centrifuged at 3,500 x g for 10 min at 4°C to isolate nucleoli. Finally, nucleoli were lysed in 300 μL RIPA buffer (150 mM NaCl, 1.0% IGEPAL, 0.5% sodium deoxycolate, 0.1% SDS, 50 mM Tris-HCl pH 8) with 0.1% protease inhibitor cocktail (Sigma-Aldrich, P8340).

### Western blotting

After electrophoresis, proteins were electrotransfered onto PVDF membranes, blocked with 5% dried skimmed milk in TBS-T (50 mM Tris pH 8.4, 0.9% NaCl, 0.05% Tween-20) for 1 h. The membranes were incubated overnight at 4°C in the presence of indicated primary antibodies in 5% skim milk. After washing with TBS-T, membranes were incubated with secondary antibodies in blocking buffer for 1 h and immunoreactive bands were detected (ECL, Bio-Rad).

### Immunoprecipitation of nucleolar MMP-2

Nucleolar lysate (150 µg) was incubated with anti-MMP-2 (1:200, ab92536) overnight at 4°C. 20 µL of 50% slurry protein G-Sepharose (ab1903259) was added and incubated for 1 h. Samples were centrifuged at 6,000 x g for 5 min and washed with wash buffer (10 mM Tris-HCl pH 7.4, 1 mM EGTA, 1 mM EDTA, 150 mM NaCl) containing 0.1% protease inhibitor cocktail. Immunoprecipated proteins were resuspended in 1x SDS loading buffer. Rabbit IgG was used as a negative control.

### Immunofluorescence microscopy

Cells were cultured on poly-L-lysine coated coverslips, fixed with 4% paraformaldehyde for 10 min, and permeabilized with ice cold methanol for 2 min. Cells were washed with PBS and blocked with 10% goat serum for 1 h at room temperature. Cells were incubated with primary antibodies overnight at 4°C. The solution was decanted and the cells were washed in PBS. Secondary antibodies were added and cells were incubated for 1 h. After washing with PBS, the cells were incubated with DAPI for 10 min. The coverslips were mounted with 50% glycerol.

To determine active sites of transcription, cells were incubated with 1 mM of 5-fluorouridine for 10 min. Cells were analyzed on a Zeiss Axioimager Z1 epifluorescence microscope using a Plan-Neofluar 40×/1.3 N.A oil immersion objective lens or a Zeiss LSM-710 laser scanning confocal microscope equipped with a 63X oil-immersion objective lens.

### Cleavage of recombinant histones

1 µg of human histones (histones H2A, H2B, H3.1, H3.2, H3.3 and H4) were incubated with 72 kDa MMP-2 activated with 4-aminophenylmercuric acetate (APMA) as described [48] Different molar ratios of histones and MMP-2 (1:1000 to 1:50) were incubated in MMP-2 buffer (150 mM NaCl, 5mM CaCl_2_, 50mM Tris-HCl pH 7.6) at 37°C for 60 min. The samples were denatured at 95°C for 5 min with loading buffer (3% SDS, 0.125 M Tris-HCl, 20% glycerol, and 0.03% bromophenol blue), electrophoresed in 16% Tris-Tricine gels and stained with Coomassie blue.

For sequencing, recombinant histone H3 cleavage products were transferred to a PVDF membrane, and stained with Coomassie blue. The bands of interest were cut and the peptide fragments sequenced using a Perkin Elmer-Applied Biosystems Procise Sequencer and Amino Acid Analyzer (Iowa State University Office of Biotechnology).

### Cleavage of nucleolar histone by endogenous MMP-2

Nucleolar lysates were resuspended in MMP-2 incubation buffer. To test whether endogenous MMP-2 activity could cleave nucleolar histone H3, the nucleolar suspension was incubated with or without 10 μM ARP-100 (MMP-2 preferring inhibitor) for 2 h at 4°C or 37°C.

### Nucleolar histone cleavage in cells

In an experiment designed to inhibit or stimulate MMP-2 activity within intact U2-OS cells, cells were incubated with 10 μM ONO-4817 (MMP-2 preferring inhibitor), or 100 μM SIN-100 (peroxynitrite generator) for 72 h. Next, cells were washed and nucleoli were isolated as described. The nucleolar lysates were resuspended in MMP-2 incubation buffer and incubated at 37°C for 4 h. A nucleolar lysate sample from vehicle treated cells were incubated at 4°C for 4 h as a negative control. To study the effect of histone acetylation on histone H3 degradation, cells were treated with 20 μM C646 (histone acetyltranferase inhibitor), or 100 ng/mL trichostatin-A (histone deacetylase inhibitor) for 24 h prior to nucleoli isolation. Histone H3 degradation was determined using anti-H3 N-terminal or C-terminal antibodies.

### Histone isolation

Histone isolation was performed using an acid extraction protocol [49] To map histone H3 cleavage site by MMP-2, isolated histones were incubated at 100:1 molar ratios with MMP-2, in presence or absence of ARP-100 (10 µM), and analyzed by immunoblotting using anti-H3 K18ac.

### Nucleolar chromatin immunoprecipitation (ChIP)-qPCR

Nucleoli were prepared from fixed U2-OS cells as previously described [31]. Briefly, cells were cross-linked by formaldehyde (0.25% in PBS for 10 min at room temperature), washed, harvested and resuspended in a total volume of 40 mL of PBS. After centrifugation, the cell pellet was resuspended in 1.0 mL of high-magnesium buffer. Nucleoli were released by sonication on ice using a Sonifier-250 (Branson Ultrasonics). Nucleoli were pelleted by centrifugation, and resuspended in 1.0 mL of low-magnesium buffer. Nucleoli were subjected to further sonication, pelleted and resuspended in 0.2 mL of 20/2 TE-buffer (pH 8.0), 2% sodium dodecyl sulfate (SDS). Following incubation at 37°C for 15 min, nucleolar structures had disappeared, as determined by microscopy, and the solution became viscous due to nucleolar DNA and further sonicated. The resulting sheared nucleolar chromatin was centrifuged, and the nucleolar chromatin supernatant was used immediately in ChIP assays.

Nucleolar chromatin supernatant (100 μL) was combined with 900 μL of PBS contained 0.1% BSA, and 20 μL of the appropriate antibody (IgG, anti-CTCF, anti-MMP-2 or anti-UBF). Following overnight incubation at 4°C, 50 μL of a 50% slurry of protein G-Dynabeads (ThermoFisher) pre-equilibrated with 0.1% BSA and 100 μg/mL E.coli RNA (ThermoFisher), and the mixture was incubated at 4°C for 4 h. Beads were recovered by centrifugation (600 x g) and washed at 4°C with 20/2 TE containing 0.2% SDS, 0.5% Triton X-100, and 150 mM NaCl; then with 20/2 TE, 0.2% SDS, 0.5% Triton X-100, and 500 mM NaCl; and with 20/2 TE-buffer. Immunoprecipitated material was eluted twice from the beads with 50 μL of 20/2 TE containing 2% SDS at 37°C for 10 min each. A total of 0.3 mL of 20/2 TE was added to the eluates, and RNAs and proteins were digested by the addition of RNAse A (5 μL of 20 mg/ml solution, ThermoFisher) followed by proteinase K (10 μL of a 14 mg/mL solution). Formaldehyde cross-links were reversed at 65°C for 4 h. DNA isolation was performed with PCR-cleanup columns (Qiagen) as per manufacturer’s instructions. qPCR was carried out on an ABI-7900HT instrument (ThermoFisher) in duplicate, with SYBR Green. The primers mapping h0, h4, h8, h18, h37.9 and h42.1 loci of human rDNA are listed in supplementary Table S1.

### MMP-2 knockdown U2-OS cells using siRNA and cell proliferation assay

The transfections mixture were prepared using MMP-2 Silencer-Select Validated siRNA sequences (ThermoFisher), siMMP-2_A_ (ID: s8852) or siMMP-2_B_ (ID: s8851), and Silencer-Selected Negative Control (Cat#: 4390843). For each 60 mm dish of cells, transfection was performed with 200 pmol siRNA using RNAiMAX (Invitrogen) as per manufacture’s instructions. After 24 h, cells were trypsinized and reseeded at 200,000 cell/dish. Separate dishes containing cells treated with siControl, siMMP-2_A_ or siMMP-2_B_ were counted at 24, 48 and 72 h.

### qPCR

RNA extraction was performed using RNeasy mini-kit (Qiagen). RNA concentration and purity ratios were analyzed using a NanoDrop-8000 spectrophotometer (ThermoScientific). 1 μg of total RNA was converted to cDNA using qScript-cDNA Supermix (Quantabio). qPCR was carried out on a LightCycler-480 instrument (Roche) in triplicate with SYBR Green (Roche). The sequence of the primers used were: MMP-2 forward, 5’-TGCTGAAGGACACACTAAAGAA-3’; MMP-2 reverse, 5’-CGCATGGTCTCGATGGTATT-3’; for pre-rRNA 5’ETS forward, 5’-GCCTTCTCTAGGGATCTGAGAG-3’; reverse, 5’-CCATAACGGAGGCAGAGACA-3’; HPRT1 forward, 5’-CTGCGATGGTGGCGTTTTTG-3’; HPRT1 reverse, 5’-ACAGCGTGTCAGCAATAACC-’3.

### Generation of MMP-2 knockout U2-OS cells

Guide RNAs used to target human MMP-2 (Supplementary Table S2) were prepared as previously described [50] using Alt-R CRISPR/Cas9 tracrRNA, ATTO-550 (IDT, Iowa) and Alt-R CRISPR/Cas9 crRNA (IDT, Iowa). Cas9 was assembled into ribonucleoprotein and transfected into U2-OS cells using lipofectamine CRISPRMAX according to the manufacturer’s instructions. 24 h following transfection, cells positive for ATTO-550 were sorted on a FACSAria-III by the Flow Cytometry Core at the University of Alberta. Cells were sorted into a 96-well plate at a density of one cell per well. Growth was monitored for 2-3 weeks until 50% confluency was reached. After sufficient expansion, ∼1×10^5^ cells were lysed in DirectPCR Lysis Reagent (Viagen Biotech) supplemented with 1 mg/ml proteinase K (ThermoFisher) overnight at 55°C, with the remaining cells transferred to a 24-well plate for continued growth. After ∼16 h, the sample was heated for 45 min at 85°C to inactivate proteinase K. 1 µL of lysate was used as a template for PCR amplification of the Cas9-target site. The resulting PCR product was sent for Sanger sequencing which confirmed Cas9-mediated gene disruption. In rescue experiments, CRISPR/Cas9 MMP-2 knockout cells were transfected with 1 μg pcDNA3 encoding 72 kDa MMP-2 [21] using Effectene reagent (Qiagen).

### Cell viability assay

Cell viability was measured using 3-(4,5-dimethylthiazole-2-yl)-2,5-diphenyltetrazolium bromide (Sigma-Aldrich), as per the manufacturer’s protocol.

### Gelatin zymography for MMP-2 activity

In brief, nucleoli were isolated and nucleolar lysates were prepared and mixed with non-reducing sample loading buffer (without DTT) and loaded onto 8% poly-acrylamide gels co-polymerized with 2 mg/mL gelatin. Following electrophoresis, gels were rinsed in 2.5% Triton X-100 (3×20 min) and kept in incubation buffer for 24 h at 37°C. Gels then were stained in 0.05% Coomassie blue for 2 h and destained with methanol.

## Statistical analysis

Results are expressed as mean ± S.E.M. The statistical significance of differences between the mean values was analyzed by Student’s t-test, or ANOVA followed by Dunnett’s post-hoc test, using GraphPad Prism 6 as indicated. Differences were considered significant at *p* < 0.05.

## Supporting information

N-terminal-sequencing-suppl-data

## Acknowledgments

This work was supported by grants from the Canadian Institutes of Health Research (FDN 143299 to RS; MOP 119515 to MH and PS-391979 to BPH), and the Natural Sciences and Engineering Research Council (RGPIN-2016-06381 to BPH).

## Notes

**Disclosure:** no conflict of interest

## References

1. Whitfield ML, George LK, Grant GD, Perou CM (2006) Common markers of proliferation. Nat Rev Cancer 6: 99–106

2. Lam YW, Trinkle-Mulcahy L, Lamond AI (2005) The nucleolus. J Cell Sci 118: 1335–1337

3. Olson MO (2004) Sensing cellular stress: another new function for the nucleolus? Sci STKE 2004: pe10

4. Allis CD, Jenuwein T (2016) The molecular hallmarks of epigenetic control. Nat Rev Genet 17: 487–500

5. Bannister AJ, Kouzarides T (2011) Regulation of chromatin by histone modifications. Cell Res 21: 381–395

6. Dhaenens M, Glibert P, Meert P, Vossaert L, Deforce D (2015) Histone proteolysis: a proposal for categorization into ‘clipping’ and ‘degradation’. Bioessays 37: 70–79

7. Duncan EM, Muratore-Schroeder TL, Cook RG, Garcia BA, Shabanowitz J, Hunt DF, Allis CD (2008) Cathepsin L proteolytically processes histone H3 during mouse embryonic stem cell differentiation. Cell 135: 284–294

8. Santos-Rosa H, Kirmizis A, Nelson C, Bartke T, Saksouk N, Cote J, Kouzarides T (2009) Histone H3 tail clipping regulates gene expression. Nat Struct Mol Biol 16: 17–22

9. Vossaert L, Meert P, Scheerlinck E, Glibert P, Van Roy N, Heindryckx B, De Sutter P, Dhaenens M, Deforce D (2014) Identification of histone H3 clipping activity in human embryonic stem cells. Stem Cell Res 13: 123–134

10. Hansen JC (2002) Conformational dynamics of the chromatin fiber in solution: determinants, mechanisms, and functions. Annu Rev Biophys Biomol Struct 31: 361–392

11. Tse C, Hansen JC (1997) Hybrid trypsinized nucleosomal arrays: identification of multiple functional roles of the H2A/H2B and H3/H4 N-termini in chromatin fiber compaction. Biochemistry 36: 11381–11388

12. Fletcher TM, Hansen JC (1995) Core histone tail domains mediate oligonucleosome folding and nucleosomal DNA organization through distinct molecular mechanisms. J Biol Chem 270: 25359–25362

13. Tse C, Sera T, Wolffe AP, Hansen JC (1998) Disruption of higher-order folding by core histone acetylation dramatically enhances transcription of nucleosomal arrays by RNA polymerase III. Mol Cell Biol 18: 4629–4638

14. Krajewski WA, Ausio J (1996) Modulation of the higher-order folding of chromatin by deletion of histone H3 and H4 terminal domains. Biochem J 316 ( **Pt 2**): 395–400

15. Wang X, He C, Moore SC, Ausio J (2001) Effects of histone acetylation on the solubility and folding of the chromatin fiber. J Biol Chem 276: 12764–12768

16. Kandasamy AD, Chow AK, Ali MA, Schulz R (2010) Matrix metalloproteinase-2 and myocardial oxidative stress injury: beyond the matrix. Cardiovasc Res 85: 413–423

17. Kwan JA, Schulze CJ, Wang W, Leon H, Sariahmetoglu M, Sung M, Sawicka J, Sims DE, Sawicki G, Schulz R (2004) Matrix metalloproteinase-2 (MMP-2) is present in the nucleus of cardiac myocytes and is capable of cleaving poly (ADP-ribose) polymerase (PARP) in vitro. FASEB J 18: 690–692

18. Wang W, Schulze CJ, Suarez-Pinzon WL, Dyck JR, Sawicki G, Schulz R (2002) Intracellular action of matrix metalloproteinase-2 accounts for acute myocardial ischemia and reperfusion injury. Circulation 106: 1543–1549

19. Chow AK, Cena J, El-Yazbi AF, Crawford BD, Holt A, Cho WJ, Daniel EE, Schulz R (2007) Caveolin-1 inhibits matrix metalloproteinase-2 activity in the heart. J Mol Cell Cardiol 42: 896–901

20. Hughes BG, Fan X, Cho WJ, Schulz R (2014) MMP-2 is localized to the mitochondria-associated membrane of the heart. Am J Physiol Heart Circ Physiol 306: H764–70

21. Ali MA, Chow AK, Kandasamy AD, Fan X, West LJ, Crawford BD, Simmen T, Schulz R (2012) Mechanisms of cytosolic targeting of matrix metalloproteinase-2. J Cell Physiol 227: 3397–3404

22. Ali MA, Cho WJ, Hudson B, Kassiri Z, Granzier H, Schulz R (2010) Titin is a Target of Matrix Metalloproteinase-2: Implications in Myocardial Ischemia/Reperfusion Injury. Circulation 122: 2039–2047

23. Sung MM, Schulz CG, Wang W, Sawicki G, Bautista-Lopez NL, Schulz R (2007) Matrix metalloproteinase-2 degrades the cytoskeletal protein alpha-actinin in peroxynitrite mediated myocardial injury. J Mol Cell Cardiol 43: 429–436

24. Kandasamy AD, Schulz R (2009) Glycogen synthase kinase-3beta is activated by matrix metalloproteinase-2 mediated proteolysis in cardiomyoblasts. Cardiovasc Res 83: 698–706

25. Hill JW, Poddar R, Thompson JF, Rosenberg GA, Yang Y (2012) Intranuclear matrix metalloproteinases promote DNA damage and apoptosis induced by oxygen-glucose deprivation in neurons. Neuroscience 220: 277–290

26. Hadler-Olsen E, Wetting HL, Ravuri C, Omair A, Rikardsen O, Svineng G, Kanapathippillai P, Winberg JO, Uhlin-Hansen L (2011) Organ specific regulation of tumour invasiveness and gelatinolytic activity at the invasive front. Eur J Cancer 47: 305–315

27. Kim K, Punj V, Kim JM, Lee S, Ulmer TS, Lu W, Rice JC, An W (2016) MMP-9 facilitates selective proteolysis of the histone H3 tail at genes necessary for proficient osteoclastogenesis. Genes Dev 30: 208–219

28. Baghirova S, Hughes BG, Poirier M, Kondo MY, Schulz R (2016) Nuclear matrix metalloproteinase-2 in the cardiomyocyte and the ischemic-reperfused heart. J Mol Cell Cardiol 94: 153–161

29. Sinha SK, Asotra K, Uzui H, Nagwani S, Mishra V, Rajavashisth TB (2014) Nuclear localization of catalytically active MMP-2 in endothelial cells and neurons. Am J Transl Res 6: 155–162

30. McManus KJ, Stephens DA, Adams NM, Islam SA, Freemont PS, Hendzel MJ (2006) The transcriptional regulator CBP has defined spatial associations within interphase nuclei. PLoS Comput Biol 2: e139

31. Murano K, Okuwaki M, Hisaoka M, Nagata K (2008) Transcription regulation of the rRNA gene by a multifunctional nucleolar protein, B23/nucleophosmin, through its histone chaperone activity. Mol Cell Biol 28: 3114–3126

32. van de Nobelen S, Rosa-Garrido M, Leers J, Heath H, Soochit W, Joosen L, Jonkers I, Demmers J, van der Reijden M, Torrano V et al. (2010) CTCF regulates the local epigenetic state of ribosomal DNA repeats. Epigenetics Chromatin 3: 19–8935-3-19

33. O’Sullivan AC, Sullivan GJ, McStay B (2002) UBF binding in vivo is not restricted to regulatory sequences within the vertebrate ribosomal DNA repeat. Mol Cell Biol 22: 657–668

34. Hake SB, Allis CD (2006) Histone H3 variants and their potential role in indexing mammalian genomes: the “H3 barcode hypothesis”. Proc Natl Acad Sci U S A 103: 6428–6435

35. Kumar S, Ratnikov BI, Kazanov MD, Smith JW, Cieplak P (2015) CleavPredict: A Platform for Reasoning about Matrix Metalloproteinases Proteolytic Events. PLoS One 10: e0127877

36. Viappiani S, Nicolescu AC, Holt A, Sawicki G, Crawford BD, Leon H, van Mulligen T, Schulz R (2009) Activation and modulation of 72kDa matrix metalloproteinase-2 by peroxynitrite and glutathione. Biochem Pharmacol 77: 826–834

37. Maze I, Wenderski W, Noh KM, Bagot RC, Tzavaras N, Purushothaman I, Elsasser SJ, Guo Y, Ionete C, Hurd YL et al. (2015) Critical Role of Histone Turnover in Neuronal Transcription and Plasticity. Neuron 87: 77–94

38. Cauwe B, Opdenakker G (2010) Intracellular substrate cleavage: a novel dimension in the biochemistry, biology and pathology of matrix metalloproteinases. Crit Rev Biochem Mol Biol 45: 351–423

39. Luger K, Mader AW, Richmond RK, Sargent DF, Richmond TJ (1997) Crystal structure of the nucleosome core particle at 2.8 A resolution. Nature 389: 251–260

40. Stricker SH, Koferle A, Beck S (2017) From profiles to function in epigenomics. Nat Rev Genet 18: 51–66

41. Nurse NP, Jimenez-Useche I, Smith IT, Yuan C (2013) Clipping of flexible tails of histones H3 and H4 affects the structure and dynamics of the nucleosome. Biophys J 104: 1081–1088

42. Hernandez-Verdun D, Roussel P, Thiry M, Sirri V, Lafontaine DL (2010) The nucleolus: structure/function relationship in RNA metabolism. Wiley Interdiscip Rev RNA 1: 415–431

43. Rosa-Garrido M, Ceballos L, Alonso-Lecue P, Abraira C, Delgado MD, Gandarillas A (2012) A cell cycle role for the epigenetic factor CTCF-L/BORIS. PLoS One 7: e39371

44. Taha MS, Nouri K, Milroy LG, Moll JM, Herrmann C, Brunsveld L, Piekorz RP, Ahmadian MR (2014) Subcellular fractionation and localization studies reveal a direct interaction of the fragile X mental retardation protein (FMRP) with nucleolin. PLoS One 9: e91465

45. Mosig RA, Dowling O, DiFeo A, Ramirez MC, Parker IC, Abe E, Diouri J, Aqeel AA, Wylie JD, Oblander SA et al. (2007) Loss of MMP-2 disrupts skeletal and craniofacial development and results in decreased bone mineralization, joint erosion and defects in osteoblast and osteoclast growth. Hum Mol Genet 16: 1113–1123

46. Turck J, Pollock AS, Lee LK, Marti HP, Lovett DH (1996) Matrix metalloproteinase 2 (gelatinase A) regulates glomerular mesangial cell proliferation and differentiation. J Biol Chem 271: 15074–15083

47. Liang YM, Wang X, Ramalingam R, So KY, Lam YW, Li ZF (2012) Novel nucleolar isolation method reveals rapid response of human nucleolar proteomes to serum stimulation. J Proteomics 77: 521–530

48. Knauper V, Murphy G (2010) Methods for studying activation of matrix metalloproteinases. Methods Mol Biol 622: 233–243

49. Bonenfant D, Coulot M, Towbin H, Schindler P, van Oostrum J (2006) Characterization of histone H2A and H2B variants and their post-translational modifications by mass spectrometry. Mol Cell Proteomics 5: 541–552

50. Cromwell CR, Sung K, Park J, Krysler AR, Jovel J, Kim SK, Hubbard BP (2018) Incorporation of bridged nucleic acids into CRISPR RNAs improves Cas9 endonuclease specificity. Nat Commun 9: 1448–018-03927-0

