## Supplementary material for "Matrix metalloproteinase-2 mediates ribosomal RNA transcription by cleaving nucleolar histones": N-terminal-sequencing-suppl-data

[Sequence Analysis]

| | | |
|---|---|---|
| Data Acquired | : | 6/6/2017 1:42:56 PM |
| Data Processed | : | 6/7/2017 5:32:19 AM |
| Reactor | : | 2 |
| Number of Cycles | : | 21 |
| Sequence Schedule | : | C:\PPSQ\SeqProg3_PDA_BGE\PVDF9-3.sch |
| Sample Name | : | Javier Garcia, blot |
| Sample Amount(pmol) | : | 25.0 |
| Sample ID | : | 8826 |
| Operator Name | : | System Administrator |
| Data File | : | 8826_06-06-2017 |
| Start Number | : | 1 |
| Method File | : | 8826_06-06-2017.lcm |
| Batch File | : | 8826_06-06-2017.lcb |
| Data Folder Path | : | C:\LabSolutions\Data\Project1\PPSQ\8826_06-06-2017 |
| Number of Analyses | : | 13 / 20 |
| Standard File | : | C:\LabSolutions\Data\Project1\PPSQ\8826_06-06-2017\PTH-AA_06-06-2017_D01.lcd |
| Data Comment | | |

[Sequence]

Q L A T K A A R K S
A P S

[Estimated Sequence]

| | 1 | 2 | 3 | 4 | 5 | 6 | 7 | 8 | 9 | 10 |
|---|---|---|---|---|---|---|---|---|---|---|
| 1st | Q | L | A | T | K | A | A | R | K | S |
| 2nd | E | G | T | K | Q | G | Q | W | Q | Q |
| 3rd | A | R | P | H | N | L | N | K | L | H |
| 4th | S | P | N | R | D | I | G | V | W | G |
| Reliability(%) | 47.4 | 40.4 | 92.4 | 100.0 | 29.2 | 100.0 | 100.0 | 100.0 | 41.2 | 68.3 |

| | 11 | 12 | 13 |
|---|---|---|---|
| 1st | A | P | S |
| 2nd | T | G | Q |
| 3rd | Q | Q | T |
| 4th | Y | Y | L |
| Reliability(%) | 87.6 | 8.9 | 9.6 |

[Evaluated Value]

| | 1 | 2 | 3 | 4 | 5 | 6 | 7 | 8 | 9 | 10 |
|---|---|---|---|---|---|---|---|---|---|---|
| D | 0.95 | 9.11 | 0.81 | 0.53 | 7.66 | 0.78 | 0.93 | 0.80 | 0.90 | 0.88 |
| E | 106.91 | 0.73 | 0.94 | 0.77 | 7.01 | 0.80 | 0.94 | 0.53 | 0.92 | 0.93 |
| N | 0.76 | 9.97 | 11.82 | 0.67 | 8.83 | 0.82 | 8.75 | 0.75 | 0.88 | 0.94 |
| Q | 525.97 | 0.22 | 0.73 | 0.76 | 26.73 | 0.77 | 22.45 | 0.73 | 20.06 | 7.52 |
| S | 45.26 | 0.58 | 0.64 | 0.64 | 5.73 | 0.87 | 0.79 | 0.74 | 0.77 | 45.44 |

| | | | | | | | | | | |
|---|---|---|---|---|---|---|---|---|---|---|
| T | 13.90 | 0.74 | 45.08 | 163.32 | 0.25 | 2.89 | 0.72 | 0.80 | 0.96 | 0.93 |
| H | 0.70 | 0.12 | 0.96 | 1.00 | 0.52 | 2.45 | 1.07 | 0.65 | 0.74 | 3.38 |
| G | 0.86 | 59.73 | 0.86 | 0.73 | 0.90 | 9.10 | 6.06 | 0.75 | 0.99 | 0.98 |
| A | 49.37 | 0.72 | 399.91 | 0.20 | 0.88 | 187.15 | 405.76 | 0.38 | 0.74 | 0.88 |
| Y | 0.91 | 4.70 | 0.86 | 0.68 | 5.52 | 2.40 | 3.90 | 0.76 | 0.86 | 0.82 |
| R | 0.50 | 29.85 | 0.59 | 0.88 | 1.08 | 0.95 | 0.97 | 62.22 | 0.48 | 0.82 |
| M | 0.79 | 3.86 | 0.72 | 0.76 | 1.48 | 0.78 | 2.64 | 0.71 | 0.91 | 0.82 |
| V | 0.78 | 15.03 | 3.06 | 0.69 | 3.66 | 2.12 | 0.87 | 0.88 | 0.84 | 0.80 |
| P | 0.54 | 20.97 | 18.52 | 0.67 | 2.20 | 0.94 | 0.88 | 0.81 | 0.90 | 0.79 |
| W | 16.93 | 0.26 | 0.56 | 0.70 | 0.73 | 0.92 | 0.97 | 0.92 | 1.19 | 0.75 |
| F | 0.67 | 9.06 | 3.54 | 0.74 | 2.02 | 1.33 | 0.90 | 0.79 | 0.92 | 0.88 |
| K | 0.71 | 3.73 | 4.66 | 9.14 | 75.12 | 0.21 | 0.56 | 0.90 | 76.73 | 0.28 |
| I | 0.72 | 10.29 | 1.51 | 0.66 | 0.82 | 3.18 | 4.45 | 0.87 | 0.79 | 0.92 |
| L | 0.14 | 243.44 | 0.26 | 0.59 | 6.18 | 4.64 | 1.47 | 0.82 | 6.68 | 0.82 |

| | 11 | 12 | 13 |
|---|---|---|---|
| D | 0.92 | 0.72 | 0.98 |
| E | 0.96 | 0.94 | 0.98 |
| N | 0.91 | 5.54 | 1.30 |
| Q | 7.63 | 28.23 | 14.44 |
| S | 0.54 | 0.73 | 15.52 |
| T | 11.67 | 0.73 | 8.28 |
| H | 1.72 | 0.78 | 2.24 |
| G | 0.86 | 37.33 | 0.77 |
| A | 101.23 | 0.64 | 0.86 |
| Y | 2.27 | 5.59 | 1.63 |
| R | 0.94 | 0.97 | 2.65 |
| M | 0.86 | 0.78 | 1.27 |
| V | 0.86 | 0.85 | 2.26 |
| P | 0.83 | 40.64 | 0.62 |
| W | 0.62 | 2.31 | 0.74 |
| F | 0.99 | 0.85 | 0.90 |
| K | 0.59 | 0.72 | 1.29 |
| I | 0.98 | 2.57 | 0.90 |
| L | 0.86 | 0.85 | 4.10 |

[Amount Yield(pmol)]

| | 1 | 2 | 3 | 4 | 5 | 6 | 7 | 8 | 9 | 10 |
|---|---|---|---|---|---|---|---|---|---|---|
| D | 2.25 | 0.00 | 0.00 | 0.00 | 0.11 | 0.14 | 0.09 | 0.00 | 0.16 | 0.00 |
| E | 3.90 | 0.00 | 0.19 | 0.00 | 0.17 | 0.17 | 0.11 | 0.00 | 0.32 | 0.10 |
| N | 1.96 | 0.03 | 0.06 | 0.00 | 0.08 | 0.06 | 0.14 | 0.07 | 0.09 | 0.00 |
| Q | 11.63 | 0.00 | 0.00 | 0.00 | 0.30 | 0.04 | 0.11 | 0.14 | 0.25 | 0.04 |
| S | 2.75 | 0.00 | 0.00 | 0.00 | 0.06 | 0.10 | 0.07 | 0.09 | 0.05 | 0.63 |
| T | 3.33 | 0.00 | 0.41 | 1.58 | 0.00 | 0.12 | 0.00 | 0.08 | 0.06 | 0.00 |
| H | 1.99 | 0.00 | 0.00 | 0.00 | 0.06 | 0.03 | 0.00 | 0.05 | 0.09 | 0.00 |
| G | 12.97 | 1.38 | 0.67 | 0.00 | 0.00 | 0.50 | 0.90 | 0.55 | 0.74 | 0.06 |

| | | | | | | | | | | |
|---|---|---|---|---|---|---|---|---|---|---|
| A | 5.91 | 0.00 | 7.97 | 0.00 | 0.00 | 3.82 | 1.19 | 0.00 | 0.05 | 0.05 |
| Y | 5.54 | 0.00 | 0.00 | 0.00 | 0.05 | 0.11 | 0.05 | 0.09 | 0.05 | 0.00 |
| R | 2.78 | 1.89 | 0.00 | 0.00 | 0.05 | 0.15 | 0.02 | 2.50 | 0.03 | 0.03 |
| M | 0.47 | 0.00 | 0.00 | 0.00 | 0.06 | 0.06 | 0.04 | 0.05 | 0.00 | 0.00 |
| V | 1.47 | 0.10 | 0.05 | 0.00 | 0.17 | 0.16 | 0.00 | 0.25 | 0.00 | 0.04 |
| P | 0.87 | 0.26 | 0.11 | 0.00 | 0.06 | 0.14 | 0.00 | 0.19 | 0.00 | 0.05 |
| W | 1.20 | 0.00 | 0.00 | 0.00 | 0.03 | 0.02 | 0.00 | 0.02 | 0.00 | 0.03 |
| F | 0.81 | 0.00 | 0.00 | 0.02 | 0.13 | 0.17 | 0.00 | 0.14 | 0.00 | 0.19 |
| K | 0.24 | 0.00 | 0.00 | 0.18 | 2.21 | 0.00 | 0.00 | 0.15 | 1.39 | 0.00 |
| I | 1.40 | 0.02 | 0.00 | 0.00 | 0.02 | 0.14 | 0.05 | 0.27 | 0.00 | 0.11 |
| L | 1.74 | 8.74 | 0.00 | 0.00 | 0.33 | 0.05 | 0.02 | 0.27 | 0.00 | 0.35 |

| | 11 | 12 | 13 |
|---|---|---|---|
| D | 0.00 | 0.43 | 0.00 |
| E | 0.03 | 0.24 | 0.00 |
| N | 0.00 | 0.17 | 0.00 |
| Q | 0.00 | 0.39 | 0.00 |
| S | 0.00 | 0.00 | 0.14 |
| T | 0.15 | 0.02 | 0.00 |
| H | 0.00 | 0.04 | 0.00 |
| G | 0.00 | 1.61 | 0.00 |
| A | 1.86 | 0.00 | 0.00 |
| Y | 0.10 | 0.17 | 0.03 |
| R | 0.78 | 0.00 | 0.06 |
| M | 0.04 | 0.02 | 0.00 |
| V | 0.08 | 0.10 | 0.13 |
| P | 0.02 | 1.31 | 0.09 |
| W | 0.03 | 0.00 | 0.05 |
| F | 0.49 | 0.06 | 0.03 |
| K | 0.10 | 0.03 | 0.00 |
| I | 0.06 | 0.32 | 0.00 |
| L | 0.14 | 0.33 | 0.13 |

[Percent Yield]

| | | |
|---|---|---|
| Amino Acid | : | D,E,N,Q,S,T,H,G,A,Y,R,C,M,V,P,W,F,K,I,L |
| Initial Yield(%) | : | 42.09 |
| Repetitive Yield(%) | : | 78.27 |
| Correlation Coef. | : | -0.809 |
| Number of Data | : | 13 |

[Repetitive Yield(%)]

| | | | |
|---|---|---|---|
| S : | 61.92(10-13) | | |
| A : | 78.24( 3- 6) | 62.16( 3- 7) | 83.08( 3-11) |
| | | 31.17( 6- 7) | 86.13( 6-11) |
| | | | 111.05( 7-11) |
| K : | 88.42( 5- 9) | | |

| | |
|---|---|
| Data File | : PTH-AA_06-06-2017_D01.lcd |
| Sample Name | : PTH-AA |
| Method File | : PTH-AA_06-06-2017.lcm |
| Background Data File | : |

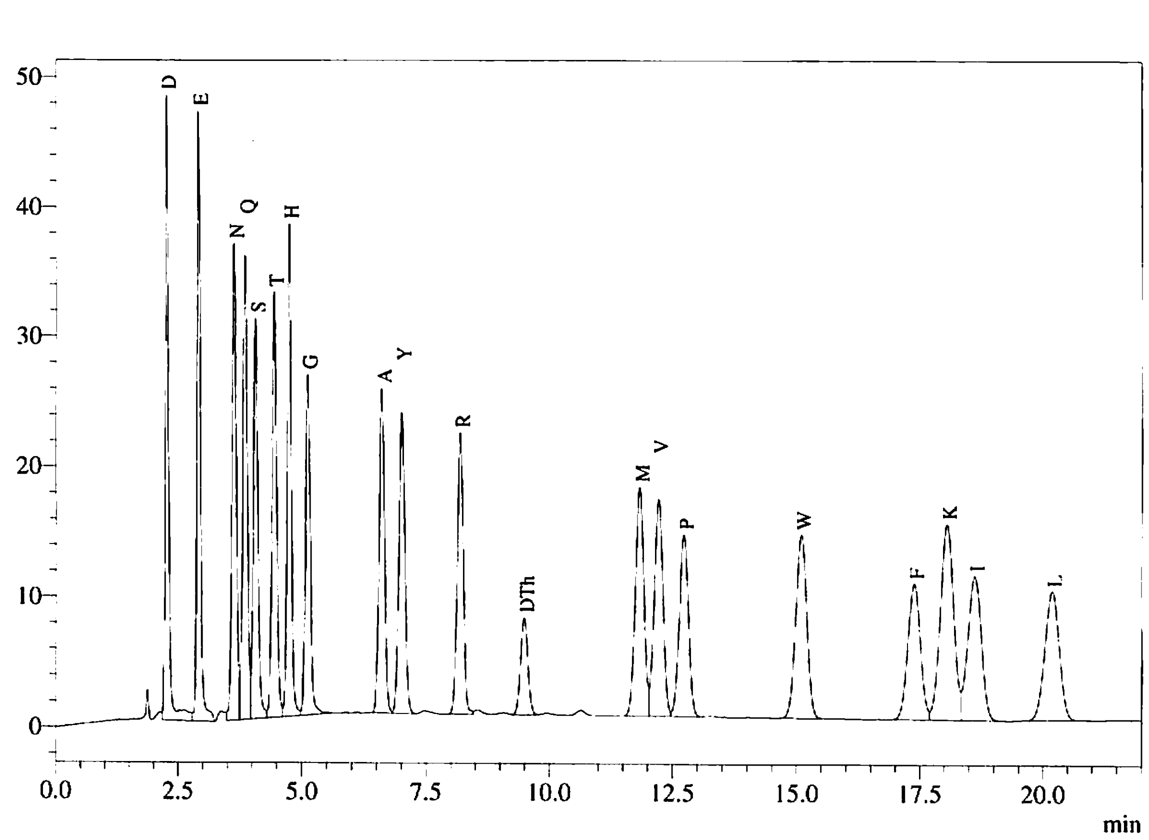


Peak Table
PDA Ch1 269nm

| Peak# | Name | Ret. Time | Area | Conc. |
|---|---|---|---|---|
| 1 | D | 2.252 | 223844 | 25.000 |
| 2 | E | 2.906 | 222572 | 25.000 |
| 3 | N | 3.625 | 222316 | 25.000 |
| 4 | Q | 3.846 | 221197 | 25.000 |
| 5 | S | 4.059 | 193581 | 25.000 |
| 6 | T | 4.437 | 211775 | 25.000 |
| 7 | H | 4.739 | 210682 | 25.000 |
| 8 | G | 5.113 | 177850 | 25.000 |
| 9 | A | 6.598 | 190336 | 25.000 |
| 10 | Y | 7.004 | 193053 | 25.000 |
| 11 | R | 8.186 | 184066 | 25.000 |
| 12 | DTh | 9.488 | 75924 | 25.000 |
| 13 | M | 11.834 | 211740 | 25.000 |
| 14 | V | 12.218 | 203748 | 25.000 |
| 15 | P | 12.727 | 177986 | 25.000 |
| 16 | W | 15.093 | 218743 | 25.000 |
| 17 | F | 17.380 | 184556 | 25.000 |
| 18 | K | 18.049 | 285299 | 25.000 |
| 19 | I | 18.610 | 207107 | 25.000 |
| 20 | L | 20.184 | 200976 | 25.000 |
| Total | | | 4017349 | |

| | |
|---|---|
| Data File | : 8826_06-06-2017_D01.lcd |
| Sample Name | : Javier Garcia, blot |
| Method File | : 8826_06-06-2017.lcm |
| Background Data File | : |

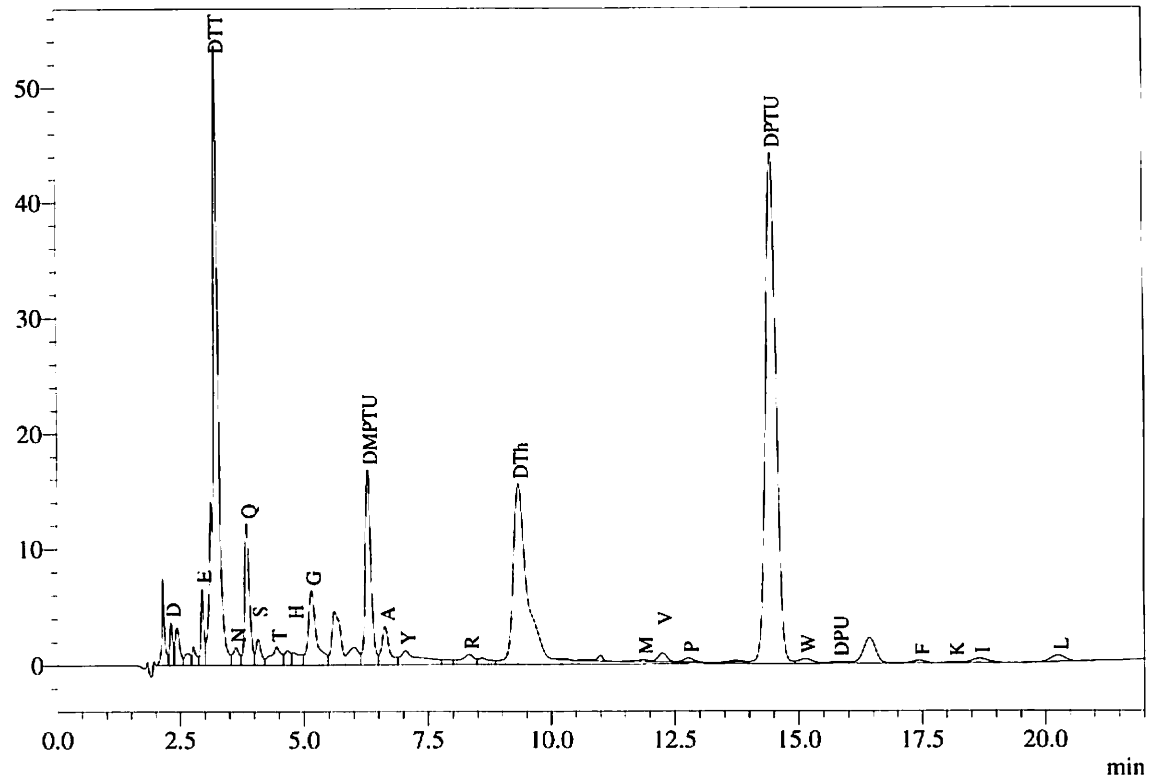


Peak Table
PDA Ch1 269nm

| Peak# | Name | Ret. Time | Area | Conc. |
|---|---|---|---|---|
| 2 | D | 2.313 | 16110 | 2.249 |
| 6 | E | 2.949 | 27770 | 3.899 |
| 7 | DTT | 3.244 | 411308 | |
| 8 | N | 3.628 | 13913 | 1.956 |
| 9 | Q | 3.855 | 82325 | 11.631 |
| 10 | S | 4.071 | 17031 | 2.749 |
| 11 | T | 4.446 | 22541 | 3.326 |
| 13 | H | 4.783 | 13440 | 1.994 |
| 14 | G | 5.152 | 73788 | 12.965 |
| 17 | DMPTU | 6.293 | 135566 | |
| 18 | A | 6.626 | 36017 | 5.913 |
| 19 | Y | 7.037 | 34240 | 5.543 |
| 21 | R | 8.322 | 16383 | 2.781 |
| 23 | DTh | 9.339 | 354306 | 145.832 |
| 28 | M | 11.864 | 3205 | 0.473 |
| 29 | V | 12.252 | 9611 | 1.474 |
| 30 | P | 12.771 | 4976 | 0.874 |
| 32 | DPTU | 14.464 | 655050 | |
| 33 | W | 15.134 | 8409 | 1.201 |
| 34 | DPU | 15.792 | 4081 | |
| 36 | F | 17.444 | 4781 | 0.810 |
| 37 | K | 18.148 | 2146 | 0.235 |
| 38 | I | 18.657 | 9284 | 1.401 |
| 40 | L | 20.259 | 11174 | 1.737 |
| Total | | | 1967455 | |

| | |
|---|---|
| Data File | : 8826_06-06-2017_D02.lcd |
| Sample Name | : Javier Garcia, blot |
| Method File | : 8826_06-06-2017.lcm |
| Background Data File | : 8826_06-06-2017_D01.lcd |

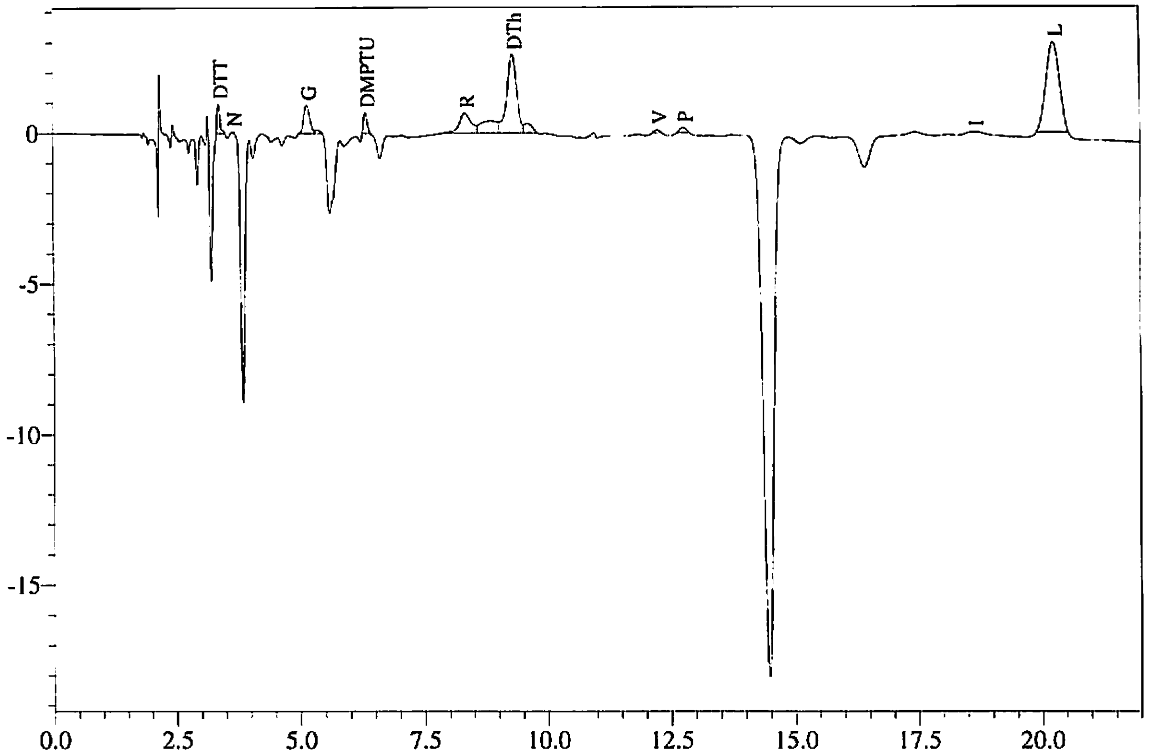


Peak Table
PDA Ch1 269nm

| Peak# | Name | Ret. Time | Area | Conc. |
|---|---|---|---|---|
| 1 | DTT | 3.370 | 4069 | |
| 2 | N | 3.654 | 214 | 0.030 |
| 3 | G | 5.151 | 7878 | 1.384 |
| 5 | DMPTU | 6.327 | 3436 | |
| 6 | R | 8.340 | 11139 | 1.891 |
| 8 | DTh | 9.309 | 36451 | 15.003 |
| 10 | V | 12.250 | 642 | 0.099 |
| 11 | P | 12.767 | 1468 | 0.258 |
| 12 | I | 18.643 | 135 | 0.020 |
| 13 | L | 20.255 | 56233 | 8.744 |
| Total | | | 121664 | |

| | |
|---|---|
| Data File | : 8826_06-06-2017_D03.lcd |
| Sample Name | : Javier Garcia, blot |
| Method File | : 8826_06-06-2017.lcm |
| Background Data File | : 8826_06-06-2017_D02.lcd |

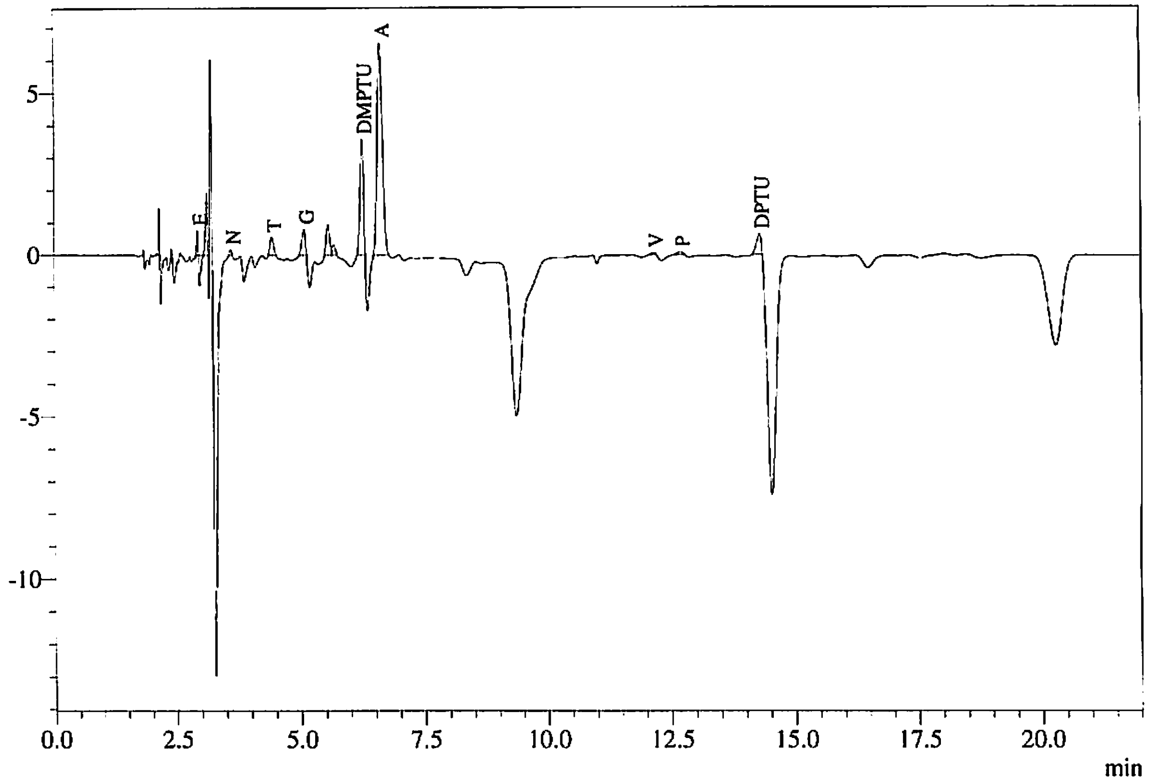


Peak Table
PDA Ch1 269nm

| Peak# | Name | Ret. Time | Area | Conc. |
|---|---|---|---|---|
| 2 | E | 2.933 | 1364 | 0.191 |
| 4 | N | 3.601 | 458 | 0.064 |
| 5 | T | 4.429 | 2747 | 0.405 |
| 6 | G | 5.080 | 3790 | 0.666 |
| 9 | DMPTU | 6.248 | 17045 | |
| 10 | A | 6.611 | 48519 | 7.966 |
| 11 | V | 12.156 | 306 | 0.047 |
| 12 | P | 12.682 | 602 | 0.106 |
| 13 | DPTU | 14.290 | 4179 | |
| Total | | | 79010 | |

| | |
|---|---|
| Data File | : 8826_06-06-2017_D04.lcd |
| Sample Name | : Javier Garcia, blot |
| Method File | : 8826_06-06-2017.lcm |
| Background Data File | : 8826_06-06-2017_D03.lcd |

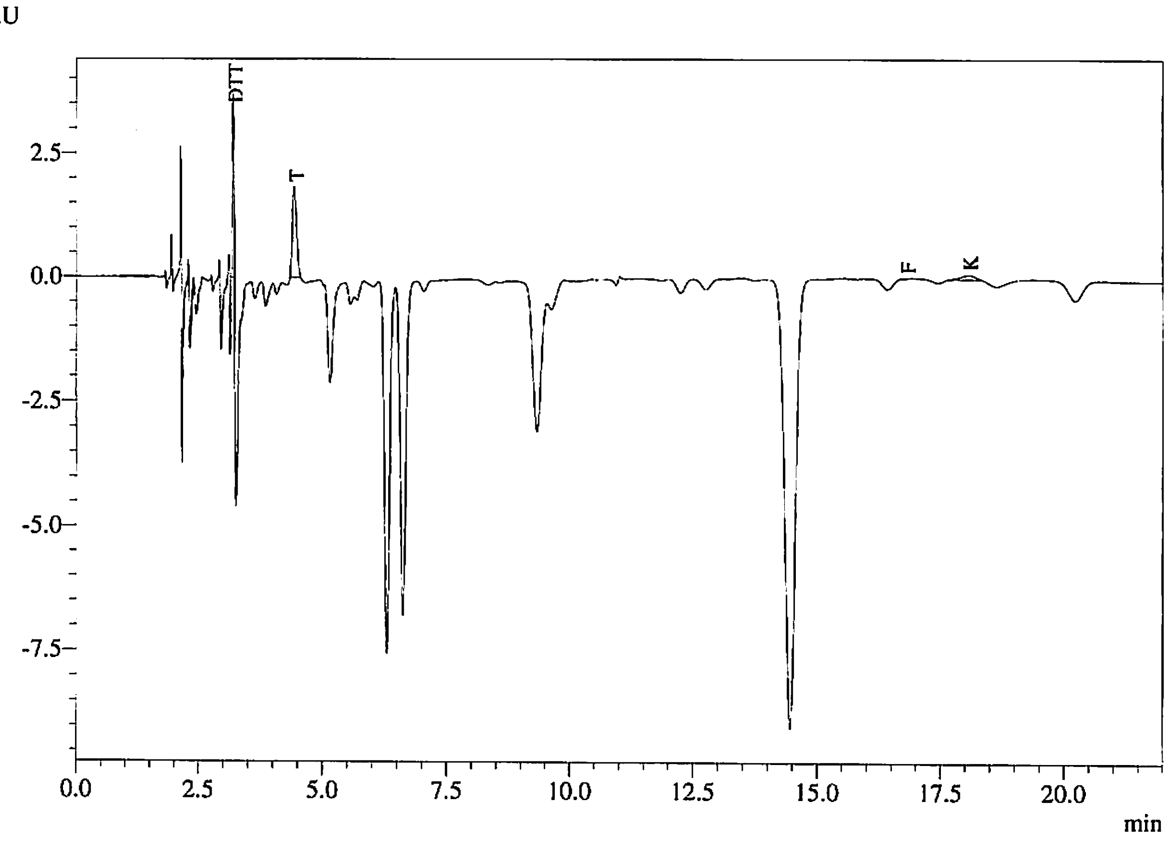


Peak Table
PDA Ch1 269nm

| Peak# | Name | Ret. Time | Area | Conc. |
|---|---|---|---|---|
| 6 | DTT | 3.196 | 7960 | |
| 7 | T | 4.429 | 10736 | 1.584 |
| 8 | F | 16.786 | 101 | 0.017 |
| 9 | K | 18.055 | 1679 | 0.184 |
| Total | | | 20476 | |

| | |
|---|---|
| Data File | : 8826_06-06-2017_D05.lcd |
| Sample Name | : Javier Garcia, blot |
| Method File | : 8826_06-06-2017.lcm |
| Background Data File | : 8826_06-06-2017_D04.lcd |

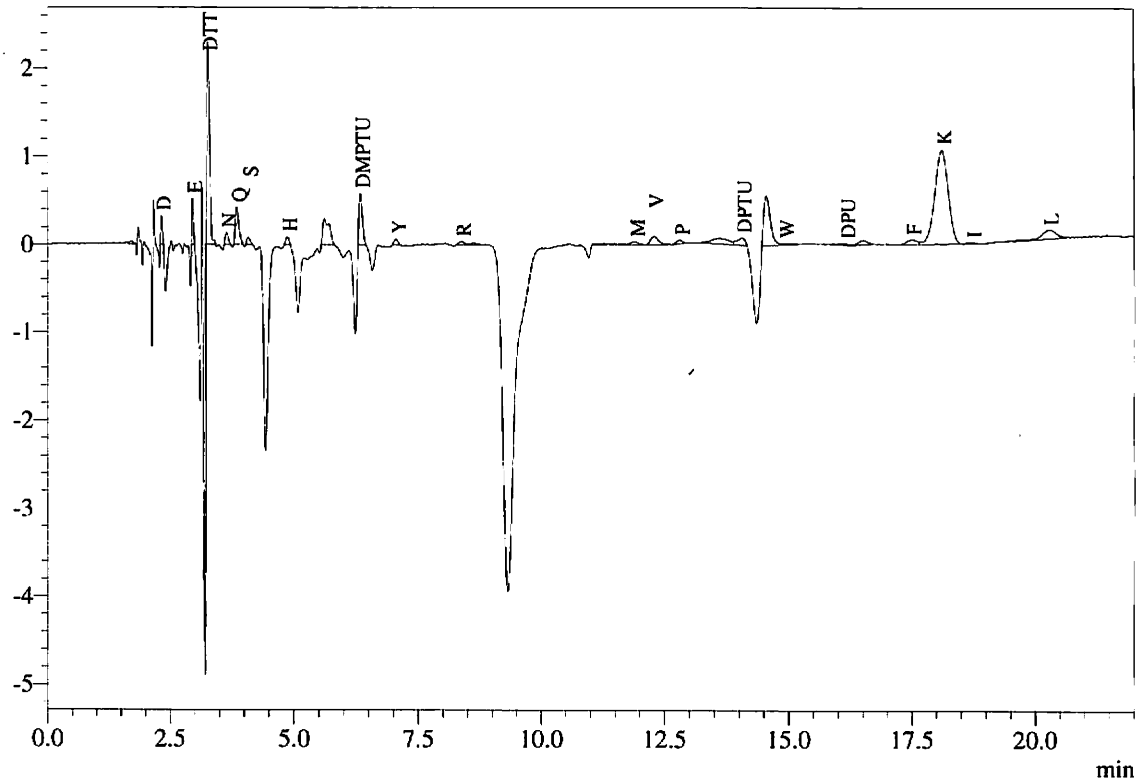


Peak Table
PDA Ch1 269nm

| Peak# | Name | Ret. Time | Area | Conc. |
|---|---|---|---|---|
| 2 | D | 2.317 | 768 | 0.107 |
| 3 | E | 2.955 | 1210 | 0.170 |
| 4 | DTT | 3.268 | 8770 | |
| 5 | N | 3.645 | 559 | 0.079 |
| 6 | Q | 3.850 | 2153 | 0.304 |
| 7 | S | 4.081 | 371 | 0.060 |
| 8 | H | 4.868 | 389 | 0.058 |
| 10 | DMPTU | 6.332 | 2733 | |
| 11 | Y | 7.049 | 330 | 0.053 |
| 12 | R | 8.372 | 319 | 0.054 |
| 18 | M | 11.915 | 430 | 0.064 |
| 19 | V | 12.293 | 1102 | 0.169 |
| 20 | P | 12.813 | 357 | 0.063 |
| 22 | DPTU | 14.062 | 927 | |
| 24 | W | 14.916 | 203 | 0.029 |
| 25 | DPU | 16.159 | 252 | |
| 27 | F | 17.498 | 783 | 0.133 |
| 28 | K | 18.112 | 20205 | 2.213 |
| 29 | I | 18.710 | 120 | 0.018 |
| 31 | L | 20.282 | 2119 | 0.329 |
| Total | | | 44101 | |

| | |
|---|---|
| Data File | : 8826_06-06-2017_D06.lcd |
| Sample Name | : Javier Garcia, blot |
| Method File | : 8826_06-06-2017.lcm |
| Background Data File | : 8826_06-06-2017_D05.lcd |

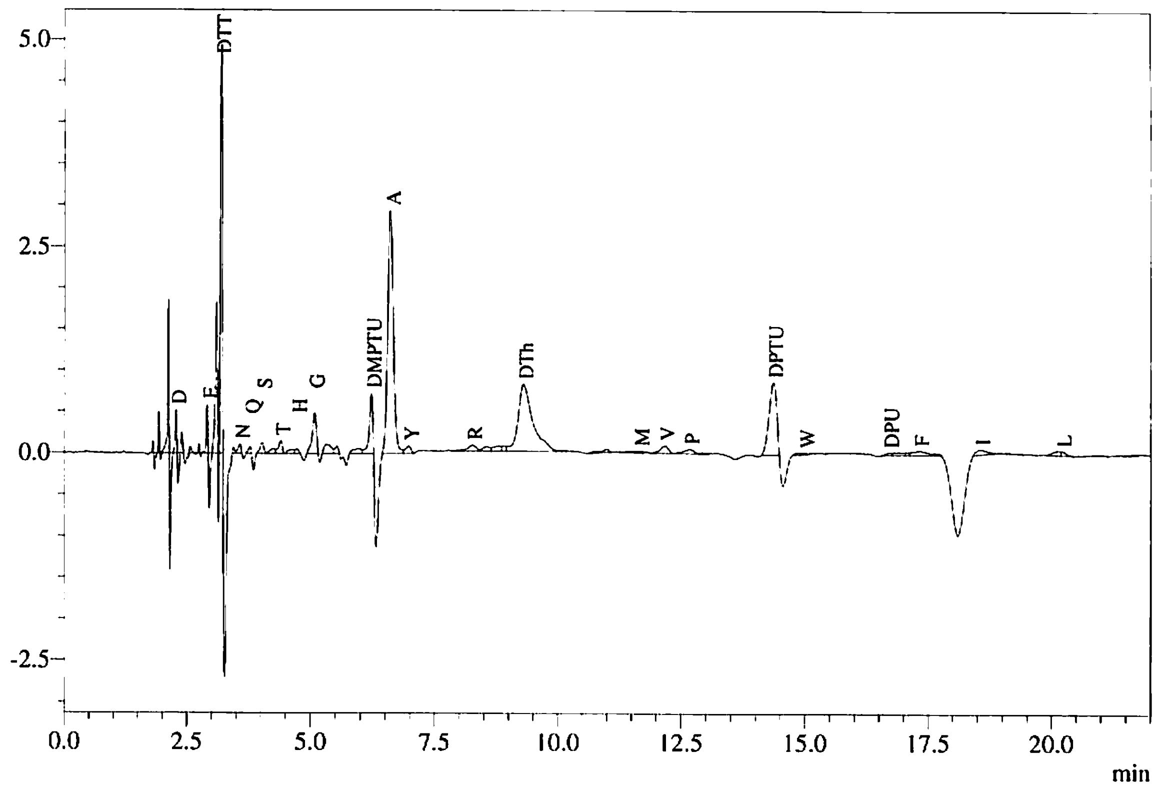


Peak Table
PDA Ch1 269nm

| Peak# | Name | Ret. Time | Area | Conc. |
|---|---|---|---|---|
| 3 | D | 2.283 | 1004 | 0.140 |
| 7 | E | 2.923 | 1177 | 0.165 |
| 9 | DTT | 3.200 | 13096 | |
| 11 | N | 3.578 | 451 | 0.063 |
| 12 | Q | 3.782 | 273 | 0.039 |
| 13 | S | 4.025 | 647 | 0.104 |
| 15 | T | 4.399 | 792 | 0.117 |
| 17 | H | 4.727 | 235 | 0.035 |
| 18 | G | 5.083 | 2832 | 0.498 |
| 22 | DMPTU | 6.224 | 3475 | |
| 23 | A | 6.604 | 23240 | 3.816 |
| 24 | Y | 6.976 | 673 | 0.109 |
| 26 | R | 8.257 | 884 | 0.150 |
| 30 | DTh | 9.302 | 16620 | 6.841 |
| 33 | M | 11.665 | 385 | 0.057 |
| 34 | V | 12.180 | 1038 | 0.159 |
| 35 | P | 12.674 | 800 | 0.140 |
| 37 | DPTU | 14.363 | 9171 | |
| 39 | W | 14.997 | 117 | 0.017 |
| 40 | DPU | 16.704 | 235 | |
| 43 | F | 17.319 | 1009 | 0.171 |
| 45 | I | 18.562 | 943 | 0.142 |
| 47 | L | 20.203 | 325 | 0.051 |
| Total | | | 79423 | |

Data File : 8826_06-06-2017_D07.lcd
Sample Name : Javier Garcia, blot
Method File : 8826_06-06-2017.lcm
Background Data File : 8826_06-06-2017_D06.lcd

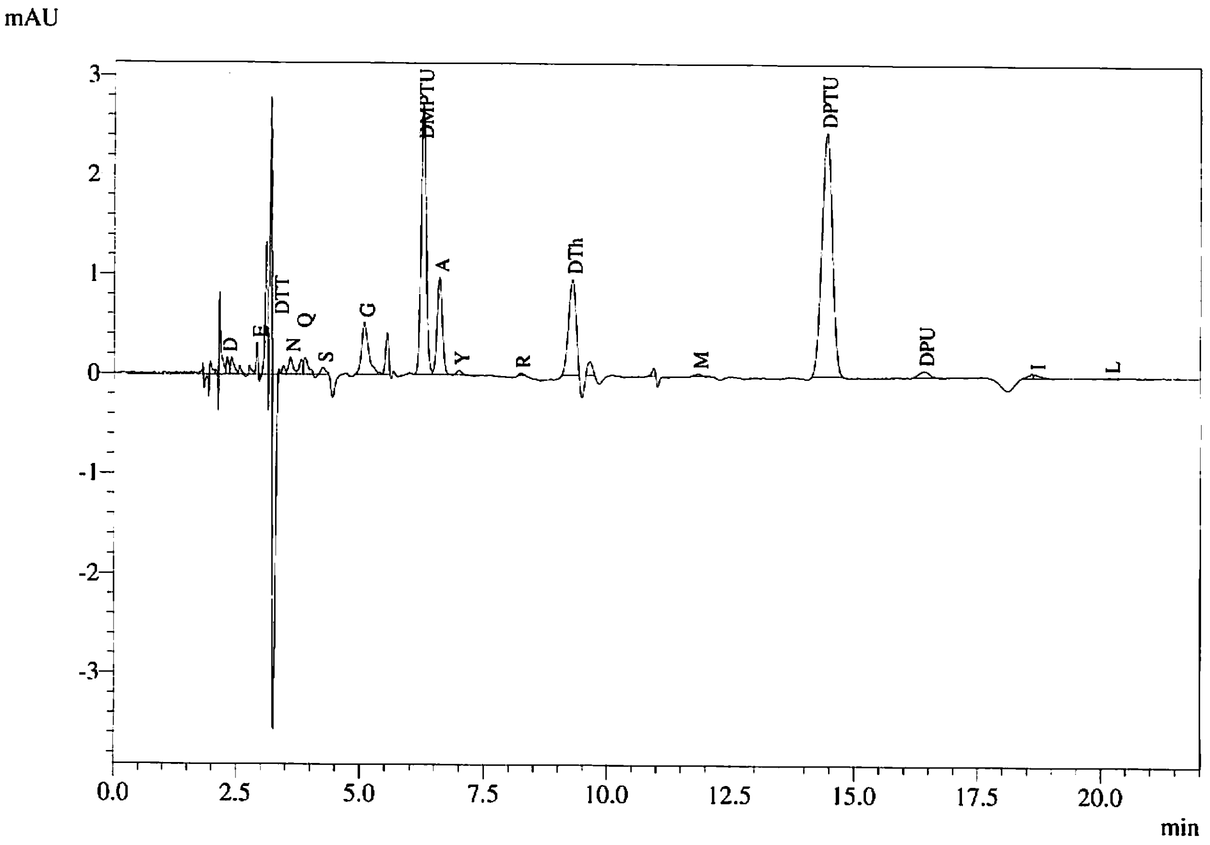


Peak Table
PDA Ch1 269nm

| Peak# | Name | Ret. Time | Area | Conc. |
|---|---|---|---|---|
| 11 | D | 2.305 | 609 | 0.085 |
| 15 | E | 2.914 | 757 | 0.106 |
| 18 | DTT | 3.360 | 113 | |
| 20 | N | 3.591 | 1022 | 0.144 |
| 21 | Q | 3.819 | 771 | 0.109 |
| 23 | S | 4.245 | 452 | 0.073 |
| 24 | G | 5.080 | 5096 | 0.895 |
| 26 | DMPTU | 6.259 | 19293 | |
| 27 | A | 6.590 | 7244 | 1.189 |
| 28 | Y | 6.991 | 309 | 0.050 |
| 29 | R | 8.228 | 125 | 0.021 |
| 30 | DTh | 9.285 | 10163 | 4.183 |
| 33 | M | 11.838 | 237 | 0.035 |
| 34 | DPTU | 14.431 | 36083 | |
| 35 | DPU | 16.399 | 974 | |
| 37 | I | 18.667 | 361 | 0.054 |
| 38 | L | 20.169 | 134 | 0.021 |
| Total | | | 83743 | |

| | |
|---|---|
| Data File | : 8826_06-06-2017_D08.lcd |
| Sample Name | : Javier Garcia, blot |
| Method File | : 8826_06-06-2017.lcm |
| Background Data File | : 8826_06-06-2017_D07.lcd |

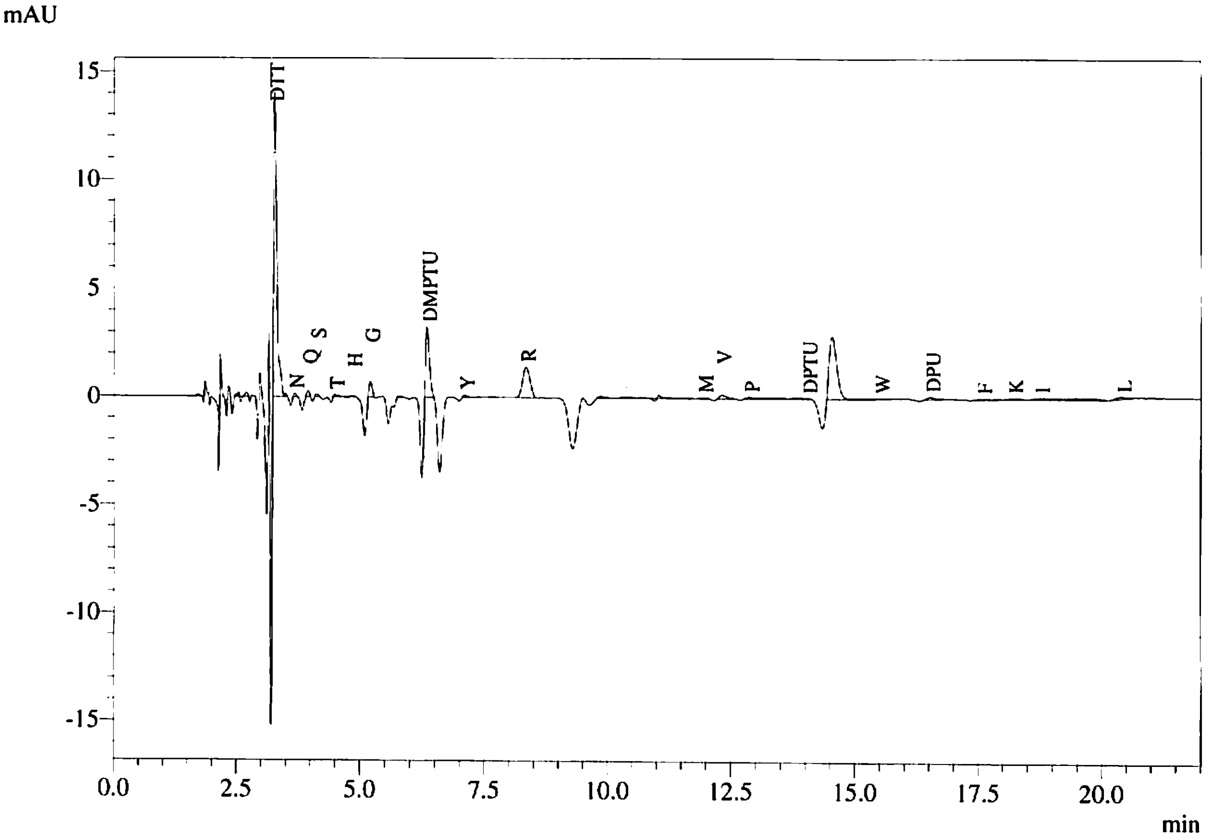


Peak Table
PDA Ch1 269nm

| Peak# | Name | Ret. Time | Area | Conc. |
|---|---|---|---|---|
| 5 | DTT | 3.267 | 62125 | |
| 7 | N | 3.668 | 526 | 0.074 |
| 8 | Q | 3.940 | 996 | 0.141 |
| 9 | S | 4.109 | 531 | 0.086 |
| 10 | T | 4.479 | 538 | 0.079 |
| 11 | H | 4.835 | 330 | 0.049 |
| 12 | G | 5.193 | 3117 | 0.548 |
| 14 | DMPTU | 6.332 | 16732 | |
| 15 | Y | 7.095 | 583 | 0.094 |
| 18 | R | 8.333 | 14751 | 2.504 |
| 25 | M | 11.933 | 329 | 0.049 |
| 26 | V | 12.313 | 1649 | 0.253 |
| 27 | P | 12.861 | 1086 | 0.191 |
| 33 | DPTU | 14.027 | 167 | |
| 35 | W | 15.507 | 123 | 0.018 |
| 36 | DPU | 16.525 | 1199 | |
| 37 | F | 17.564 | 798 | 0.135 |
| 39 | K | 18.210 | 1359 | 0.149 |
| 40 | I | 18.745 | 1757 | 0.265 |
| 48 | L | 20.406 | 1761 | 0.274 |
| Total | | | 110457 | |

Data File : 8826_06-06-2017_D09.lcd
Sample Name : Javier Garcia, blot
Method File : 8826_06-06-2017.lcm
Background Data File : 8826_06-06-2017_D08.lcd

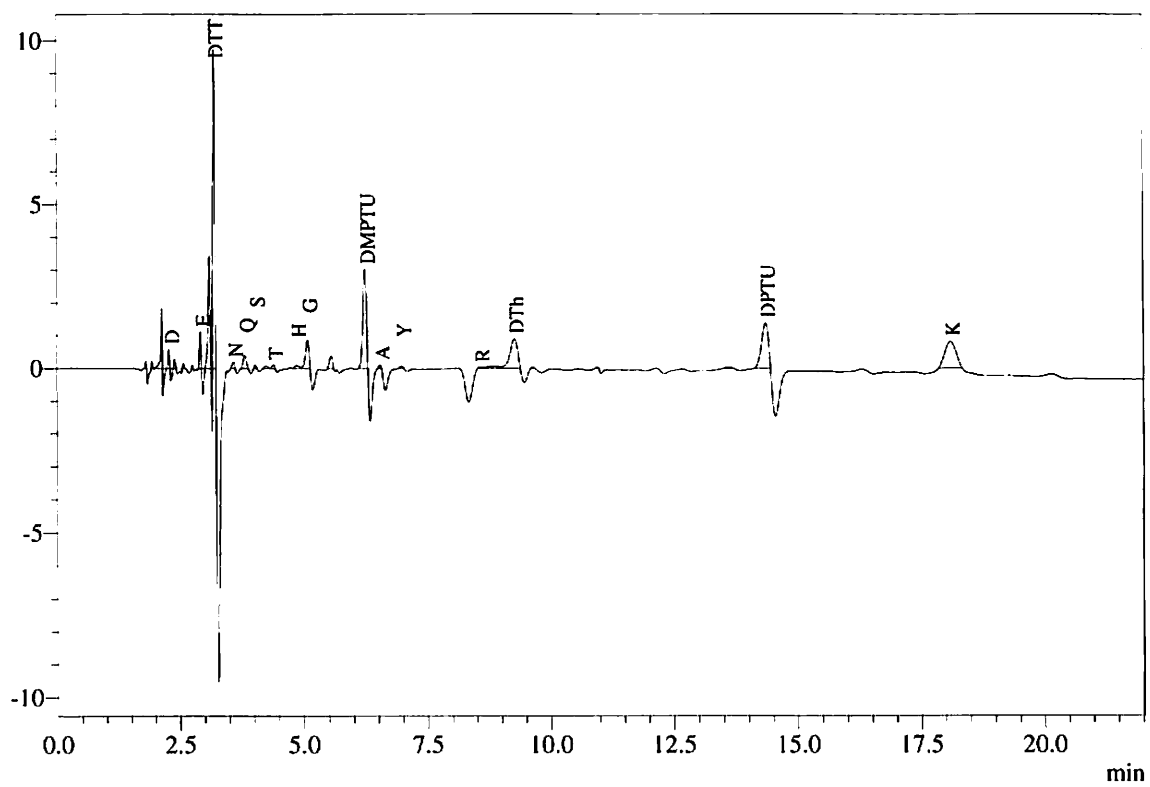


Peak Table
PDA Ch1 269nm

| Peak# | Name | Ret. Time | Area | Conc. |
|---|---|---|---|---|
| 2 | D | 2.293 | 1128 | 0.158 |
| 5 | E | 2.933 | 2309 | 0.324 |
| 7 | DTT | 3.206 | 23127 | |
| 8 | N | 3.592 | 628 | 0.088 |
| 9 | Q | 3.820 | 1754 | 0.248 |
| 10 | S | 4.034 | 303 | 0.049 |
| 12 | T | 4.402 | 378 | 0.056 |
| 13 | H | 4.867 | 634 | 0.094 |
| 14 | G | 5.090 | 4192 | 0.737 |
| 16 | DMPTU | 6.245 | 14502 | |
| 17 | A | 6.547 | 302 | 0.050 |
| 18 | Y | 6.981 | 296 | 0.048 |
| 19 | R | 8.568 | 171 | 0.029 |
| 21 | DTh | 9.267 | 9614 | 3.957 |
| 25 | DPTU | 14.356 | 12987 | |
| 26 | K | 18.101 | 12684 | 1.389 |
| Total | | | 85009 | |

| | |
|---|---|
| Data File | : 8826_06-06-2017_D10.lcd |
| Sample Name | : Javier Garcia, blot |
| Method File | : 8826_06-06-2017.lcm |
| Background Data File | : 8826_06-06-2017_D09.lcd |

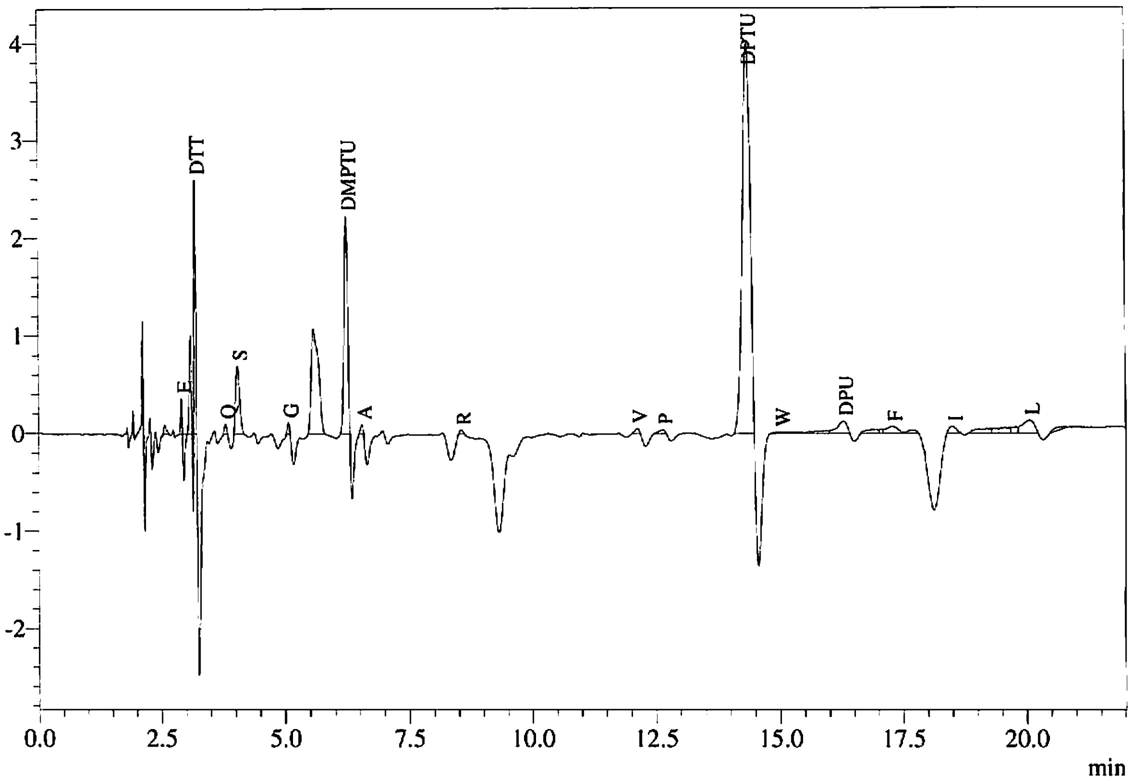


Peak Table
PDA Ch1 269nm

| Peak# | Name | Ret. Time | Area | Conc. |
|---|---|---|---|---|
| 2 | E | 2.923 | 694 | 0.097 |
| 4 | DTT | 3.197 | 5920 | |
| 5 | Q | 3.807 | 313 | 0.044 |
| 6 | S | 4.050 | 3883 | 0.627 |
| 7 | G | 5.077 | 334 | 0.059 |
| 9 | DMPTU | 6.240 | 11340 | |
| 10 | A | 6.547 | 305 | 0.050 |
| 11 | R | 8.555 | 154 | 0.026 |
| 12 | V | 12.129 | 292 | 0.045 |
| 13 | P | 12.645 | 301 | 0.053 |
| 14 | DPTU | 14.360 | 43653 | |
| 15 | W | 15.018 | 238 | 0.034 |
| 19 | DPU | 16.290 | 1710 | |
| 22 | F | 17.284 | 1108 | 0.188 |
| 24 | I | 18.497 | 708 | 0.107 |
| 29 | L | 20.060 | 2232 | 0.347 |
| Total | | | 73186 | |

| | |
|---|---|
| Data File | : 8826_06-06-2017_D11.lcd |
| Sample Name | : Javier Garcia, blot |
| Method File | : 8826_06-06-2017.lcm |
| Background Data File | : 8826_06-06-2017_D10.lcd |

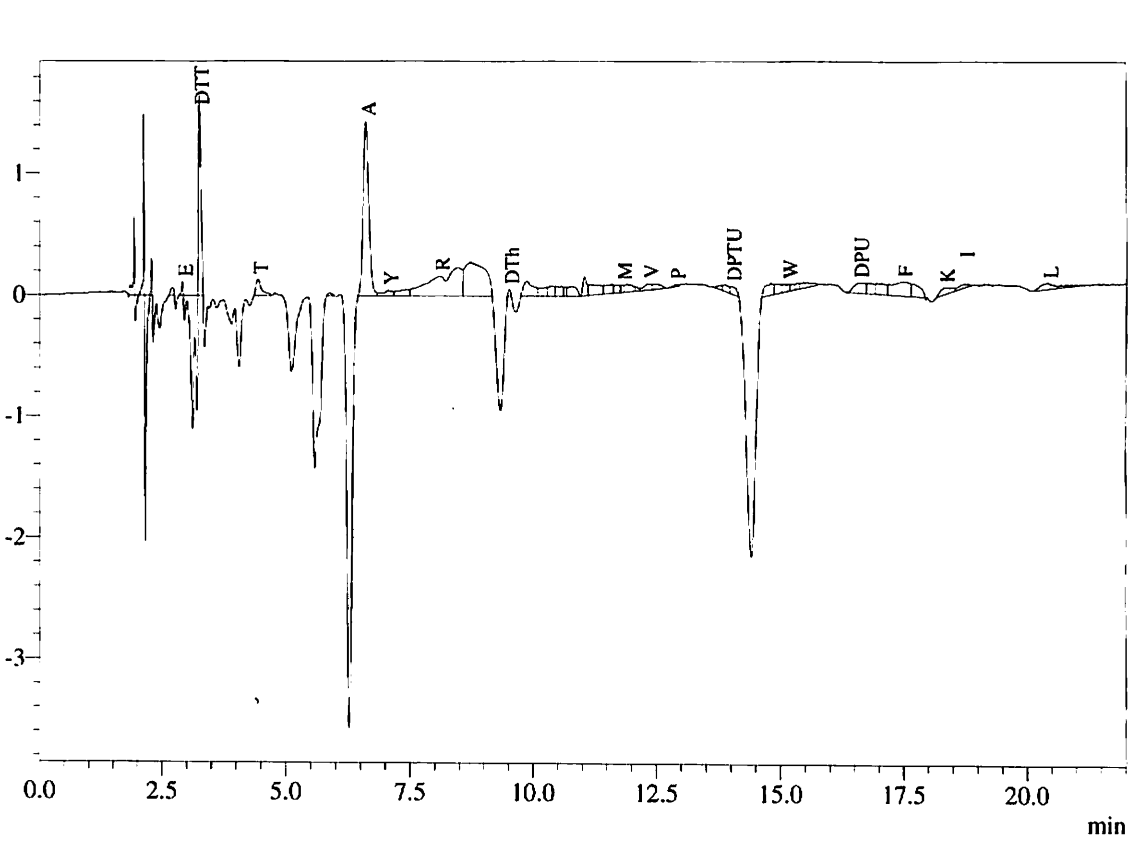


Peak Table
PDA Ch1 269nm

| Peak# | Name | Ret. Time | Area | Conc. |
|---|---|---|---|---|
| 4 | E | 2.912 | 209 | 0.029 |
| 5 | DTT | 3.258 | 5603 | |
| 6 | T | 4.440 | 1032 | 0.152 |
| 8 | A | 6.607 | 11319 | 1.858 |
| 9 | Y | 7.055 | 594 | 0.096 |
| 11 | R | 8.095 | 4604 | 0.782 |
| 14 | DTh | 9.510 | 167 | 0.069 |
| 25 | M | 11.808 | 258 | 0.038 |
| 28 | V | 12.340 | 550 | 0.084 |
| 30 | P | 12.875 | 124 | 0.022 |
| 33 | DPTU | 14.005 | 547 | |
| 37 | W | 15.147 | 215 | 0.031 |
| 41 | DPU | 16.582 | 1372 | |
| 44 | F | 17.468 | 2882 | 0.488 |
| 46 | K | 18.325 | 898 | 0.098 |
| 47 | I | 18.727 | 428 | 0.065 |
| 48 | L | 20.431 | 919 | 0.143 |
| Total | | | 31723 | |

| | |
|---|---|
| Data File | : 8826_06-06-2017_D12.lcd |
| Sample Name | : Javier Garcia, blot |
| Method File | : 8826_06-06-2017.lcm |
| Background Data File | : 8826_06-06-2017_D11.lcd |

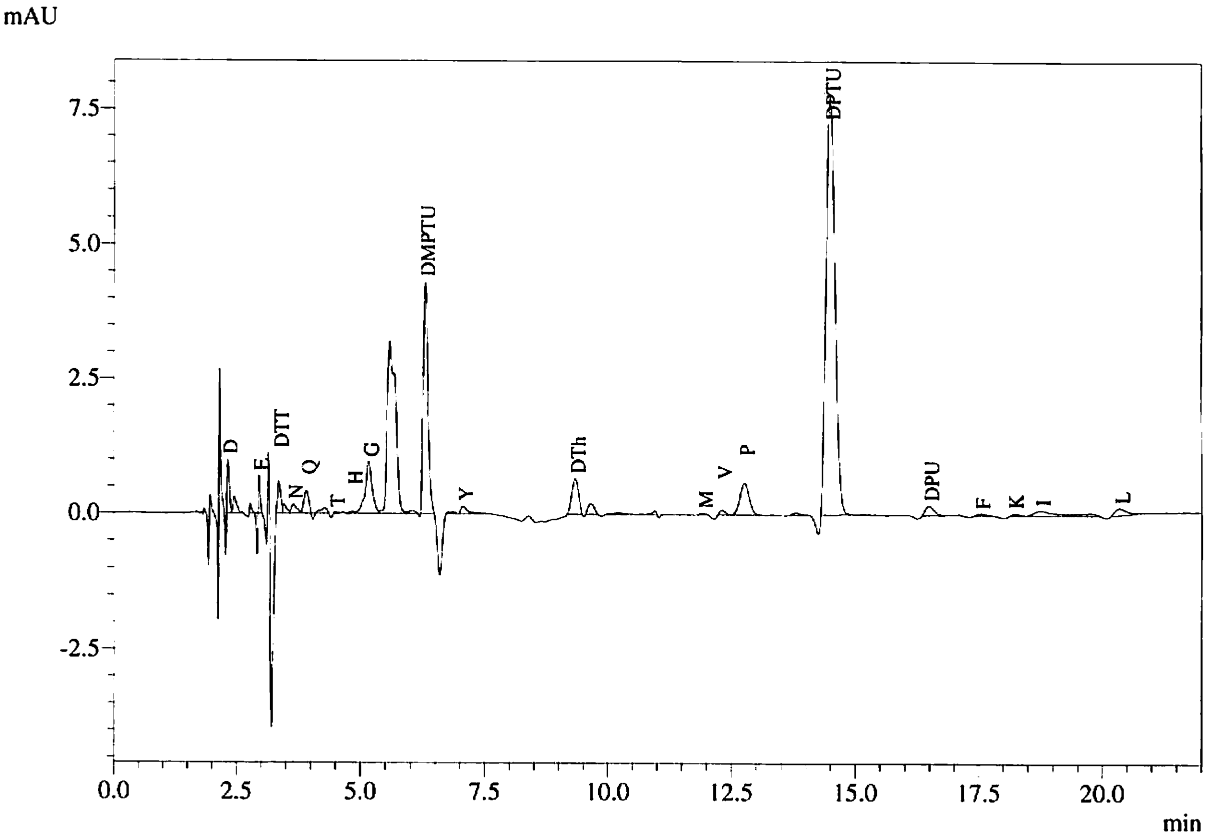


Peak Table
PDA Ch1 269nm

| Peak# | Name | Ret. Time | Area | Conc. |
|---|---|---|---|---|
| 1 | D | 2.320 | 3079 | 0.430 |
| 4 | E | 2.956 | 1678 | 0.236 |
| 5 | DTT | 3.358 | 2669 | |
| 7 | N | 3.645 | 1190 | 0.167 |
| 8 | Q | 3.913 | 2771 | 0.391 |
| 10 | T | 4.490 | 106 | 0.016 |
| 11 | H | 4.848 | 242 | 0.036 |
| 12 | G | 5.167 | 9157 | 1.609 |
| 15 | DMPTU | 6.300 | 28854 | |
| 17 | Y | 7.063 | 1035 | 0.167 |
| 19 | DTh | 9.337 | 6173 | 2.541 |
| 23 | M | 11.932 | 119 | 0.018 |
| 24 | V | 12.321 | 621 | 0.095 |
| 25 | P | 12.766 | 7446 | 1.307 |
| 27 | DPTU | 14.490 | 96909 | |
| 28 | DPU | 16.490 | 2067 | |
| 29 | F | 17.536 | 366 | 0.062 |
| 30 | K | 18.205 | 292 | 0.032 |
| 31 | I | 18.756 | 2090 | 0.315 |
| 37 | L | 20.363 | 2115 | 0.329 |
| Total | | | 168978 | |

| | |
|---|---|
| Data File | : 8826_06-06-2017_D13.lcd |
| Sample Name | : Javier Garcia, blot |
| Method File | : 8826_06-06-2017.lcm |
| Background Data File | : 8826_06-06-2017_D12.lcd |

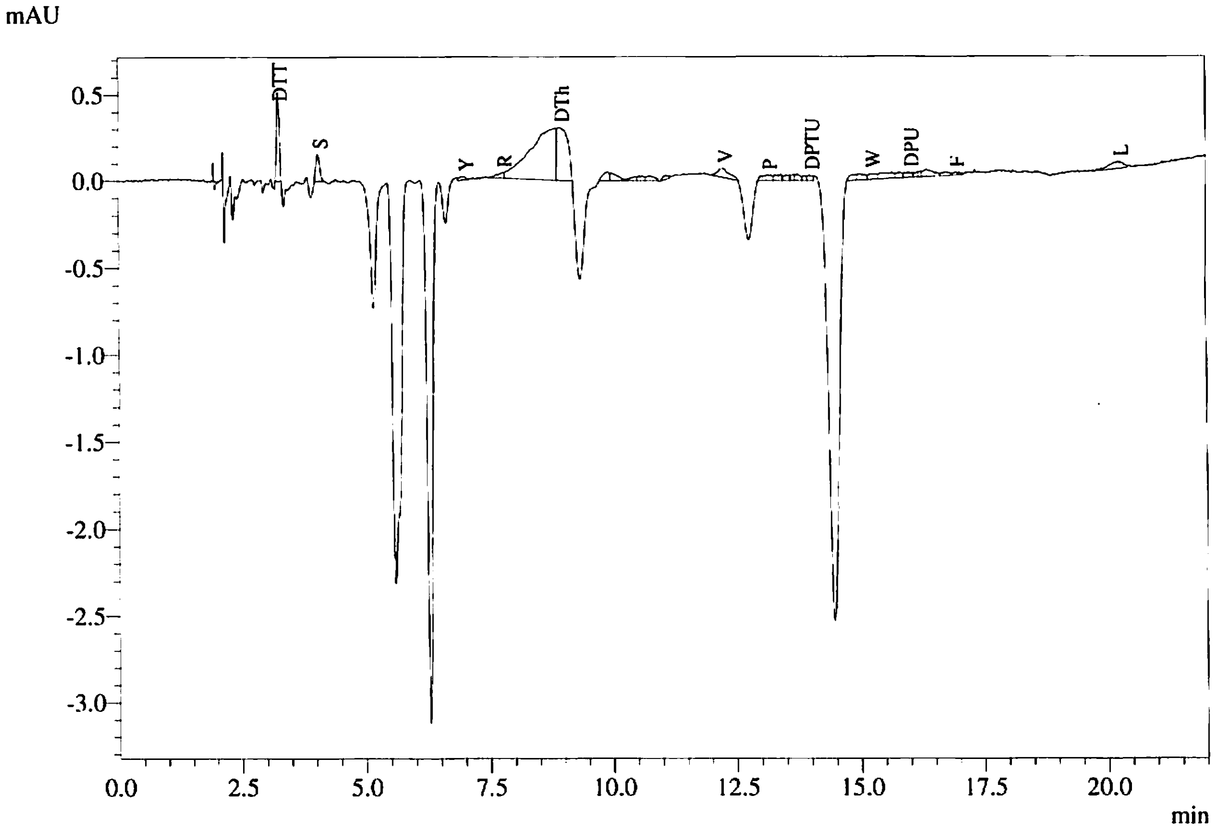


Peak Table
PDA Ch1 269nm

| Peak# | Name | Ret. Time | Area | Conc. |
|---|---|---|---|---|
| 2 | DTT | 3.252 | 2062 | |
| 3 | S | 4.058 | 850 | 0.137 |
| 4 | Y | 6.997 | 184 | 0.030 |
| 5 | R | 7.765 | 341 | 0.058 |
| 7 | DTh | 8.925 | 4732 | 1.948 |
| 14 | V | 12.247 | 873 | 0.134 |
| 15 | P | 13.143 | 502 | 0.088 |
| 21 | DPTU | 13.995 | 229 | |
| 24 | W | 15.221 | 376 | 0.054 |
| 27 | DPU | 15.991 | 331 | |
| 32 | F | 16.917 | 165 | 0.028 |
| 33 | L | 20.245 | 831 | 0.129 |
| Total | | | 11476 | |
