## Supplementary Material for "Matrix metalloproteinase-2 mediates ribosomal RNA transcription by cleaving nucleolar histones"

**Supplementary Tables**

### Table S1. Primer pairs used for ChIP assays of the rRNA gene

| **Region** | **Forward (5’-3’)** | **Reverse (5’-3’)** |
| --- | --- | --- |
| *h37.9* | CCCTGGTCGATTAGTTGTGG | GTGCTCCCTTCCTCTGTGAG |
| *h42.1* | GCTTCTCGACTCACGGTTTC | CCGAGAGCACGATCTCAAA |
| *h0* | CTGCGATGGTGGCGTTTTTG | ACAGCGTGTCAGCAATAACC |
| *h4* | CCGACGACCCATTCGAACGTCT | CTCTCCGGAATCGAACCCTGA |
| *h8* | AGTCGGGTTGCTTGGGAATGC | CCCTTACGGTACTTGTTGACT |
| *h18* | GTTGACGTACAGGGTGGACTG | GGAAGTTGTCTTCACGCCTGA |

**Table S2. CRISPR RNA sequence for MMP-2 knockout**

| **Gene** | **CRISPR RNA Sequence** |
| --- | --- |
| MMP2 | ACATCTGGGTTGCCGCAGCG |

**Supplementary Figures**

**Fig. S1. Localization of MMP-2 to nucleoli of cancer cell lines**

Staining and co-localization of MMP-2 and fibrillarin within nucleoli in MCF7 (upper), PC-3 (middle) and U-373 (lower) cells. In the merged images (to the right), MMP-2 (green) and fibrillarin (red).

**Fig. S2. Viability of U-2 OS cells at increasing SIN-1, ONO-4817, and GM6001 concentrations.**

Effect of SIN-1 **(A)**, ONO-4817 **(B)**, and GM6001 **(C)** on U-2 OS cell viability as measured by the MTT assay. Cells were incubated with the compounds for 72 h at 37°C. Data represent the mean ± SD for 3 independent experiments. Calculated SIN-1 LC_50_= 325 µM, ONO-4817 LC_50_= 71 µM, and GM6001 LC_50_= 95 µM.
