## Supplementary figures and images for "Matrix metalloproteinase-2 mediates ribosomal RNA transcription by cleaving nucleolar histones"

### Supplementary Fig S1

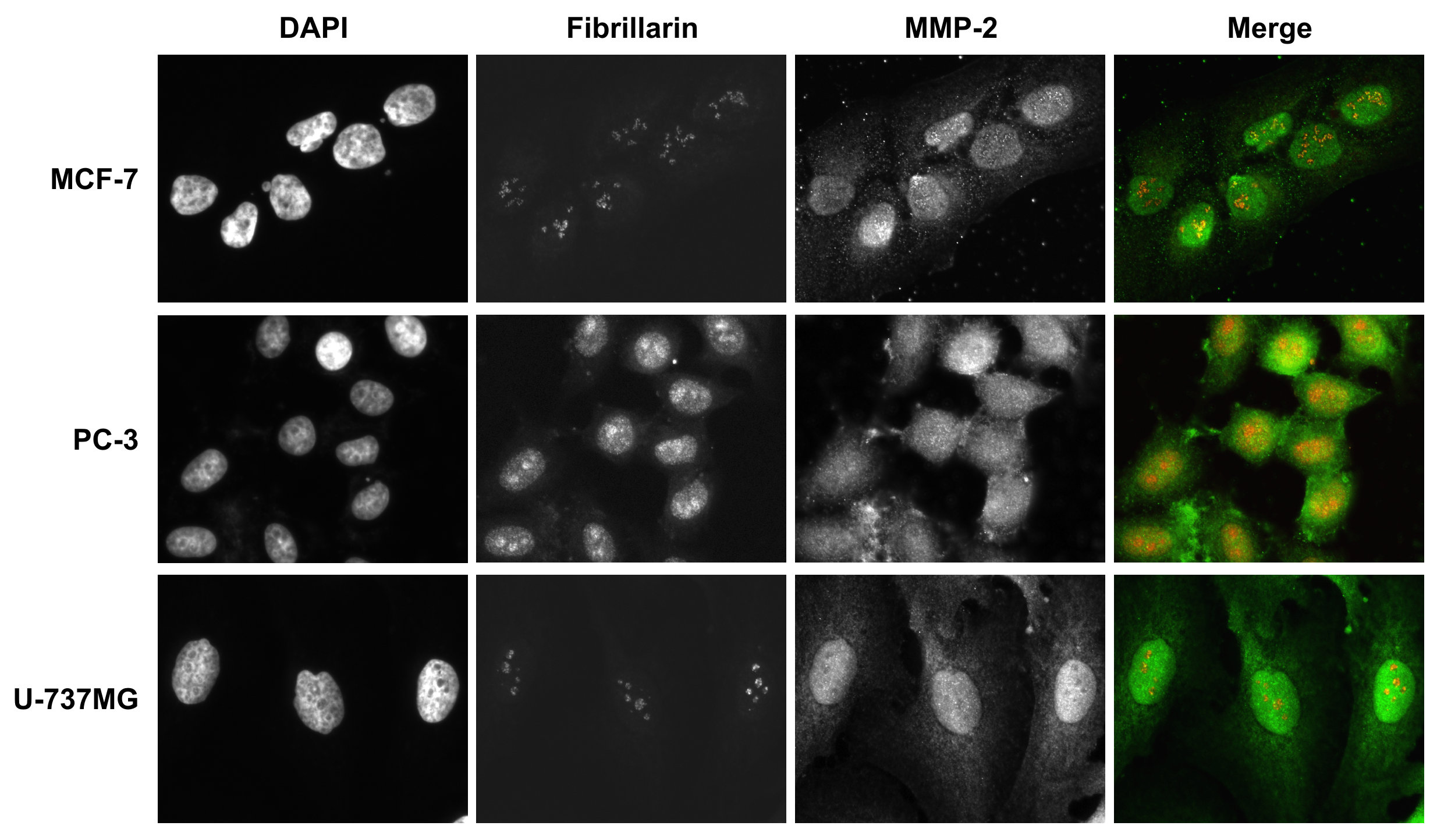

### Supplementary Fig S2

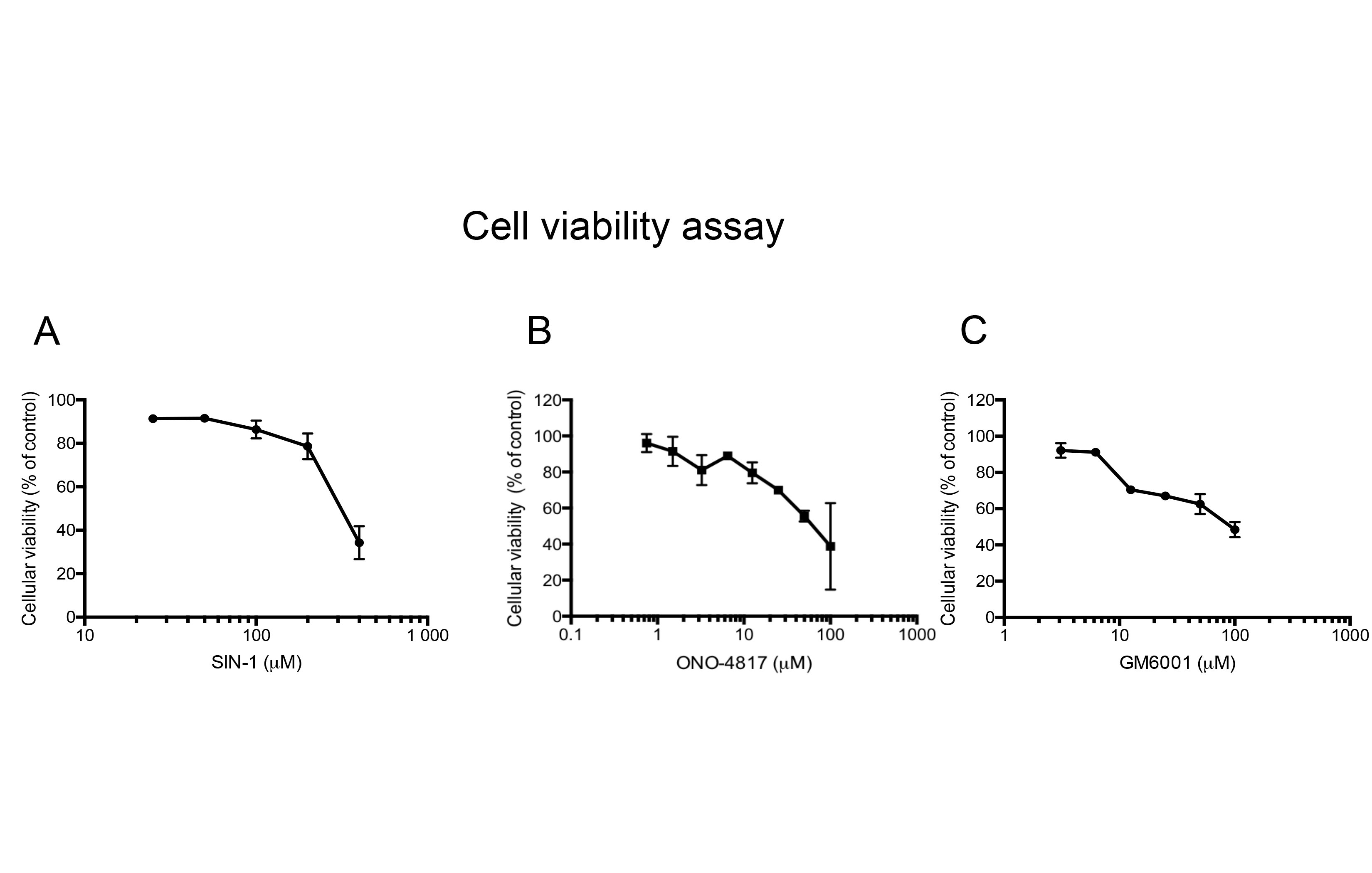
